# DHH1/DDX6-like RNA helicases maintain ephemeral half-lives of stress-response mRNAs associated with innate immunity and growth inhibition

**DOI:** 10.1101/772087

**Authors:** Thanin Chantarachot, Reed S. Sorenson, Maureen Hummel, Haiyan Ke, Alek T. Kettenburg, Daniel Chen, Karen Aiyetiwa, Katayoon Dehesh, Thomas Eulgem, Leslie E. Sieburth, Julia Bailey-Serres

## Abstract

Gene transcription is counterbalanced by mRNA decay processes that regulate transcript quality and quantity. We show here that the evolutionarily conserved DHH1/DDX6-like RNA HELICASEs of *Arabidopsis thaliana* control the ephemerality of a subset of cellular mRNAs. These RNA helicases co-localize with key markers of processing bodies and stress granules and contribute to their subcellular dynamics. These RHs function to limit the precocious accumulation and translation of stress-responsive mRNAs associated with autoimmunity and growth inhibition under non-stress conditions. Given the conservation of this RH subfamily, they may control basal levels of conditionally-regulated mRNAs in diverse eukaryotes, accelerating responses without penalty.

## Introduction

The dynamic regulation of mRNA translation, decay and sequestration is essential for growth, development and responses to internal and external stimuli. These processes involve interconnected mRNA-ribonucleoprotein (mRNP) complexes including poly(ribo)somes, processing bodies (PBs) and stress granules (SGs)^1^. Depending on the biological context, mRNAs targeted to PBs can be degraded or stabilized^2–5^, whereas those sequestered in SGs are generally stabilized^6, 7^.

In eukaryotes, the bulk of cytoplasmic mRNAs is degraded by general decay pathways that are initiated by deadenylation of the poly(A) tail. These pathways include 5’-to-3’ decay that requires mRNA decapping or 3’-to-5’ decay by the RNA exosome or the exoribonuclease SUPPRESSOR OF VARICOSE (SOV/DIS3L2). The decapping pathway requires the enzyme DECAPPING 2 (DCP2) and core decapping factors DCP1 and VARICOSE (VCS/EDC4/HEDLS) and is facilitated by conserved activators such as DCP5, PROTEIN ASSOCIATED WITH TOPOISOMERASE 1 (PAT1), the LSM1-7 complex, and DHH1/DDX6 (CGH-1/Me31B/Xp54)^8^. Decapped mRNAs can be hydrolyzed by the 5’-to-3’ exoribonuclease XRN1/4^9^. These 5’ and 3’ pathways have substrate specificity, but are not mutually exclusive. When decapping-dependent decay is compromised in Arabidopsis, the 3’-to-5’ exoribonuclease SOV can compensate to control mRNA abundance and homeostasis^10, 11^.

Spatiotemporal regulation of mRNA decay is critical for the cellular transcriptome adjustment in response to both developmental and environmental cues in plants^1^. Dysfunction in decapping due to loss-of-function of non-redundant components results in post-embryonic lethality (DCP1, DCP2, VCS, and DCP5) or severe growth alterations (LSM1 and PAT1)^12–16^. The cause of the developmental defects in some decapping mutants is associated with disruption of mRNA quality control and small interfering (si)RNA production^17^. However, there is limited knowledge of the role of the decay machinery in the spatial and temporal turnover of specific mRNAs and the connections between turnover and mRNA translation and mobilization to PBs and SGs. Mutations in the mRNA decay machinery have been identified in genetic screens for altered sensitivity to biotic and abiotic stresses^16, 18–24^, yet there is poor understanding of the importance of mRNA decay in restricting accumulation of mRNAs that provide stress resilience but constrain growth.

The DHH1/DDX6 family of DEAD-box RNA helicases is conserved across eukaryotes^25^. These proteins function at the nexus between mRNA translation, storage and decay, mediating translational repression and initiating mRNA degradation^26–33^. For example, yeast DHH1 can activate mRNA decapping^34^, promote translational repression^35^, and associate with ribosomes to sense the codon-dependent rate of translational elongation to trigger cotranslational decay^36^. However, the transcript-specific role of these helicases is generally understudied. Here we identify the Arabidopsis DHH1/DDX6-like proteins RNA HELICASE 6 (RH6), RH8, and RH12 as additive and functionally redundant mRNA decay factors required for growth and development. Severe deficiency of *RH6*, *RH8* and *RH1*2 function impairs PB and SG formation, and shifts the transcriptome and translatome homeostasis so that defense- and other stress-responsive mRNAs accumulate despite when grown under standard conditions. This coincides with a simultaneous repression of mRNAs required for general growth. RNA decay analysis determined that these RHs facilitate turnover of specific short-lived decapping substrates, enriched for stress and defense responses. The stabilization of these ephemeral mRNAs in the *rh6812* mutant confers autoimmunity. We propose that RH-mediated decay of stress-responsive mRNAs under non-stress conditions is required to maintain the growth-defense balance in plants.

## Results

### Arabidopsis RH6, RH8, and RH12 are essential DHH1/DDX6-like proteins

The *Viridiplantae* encode DHH1/DDX6-like proteins with a dual RecA helicase core, including Arabidopsis RH6 (At2g45810), RH8 (At4g00660) and RH12 (At3g61240) (Fig. 1a **and** Supplementary Fig. 1). These three RHs share 79-86% protein sequence identity and are generally co-expressed (Supplementary Fig. 2). Single gene partial and complete loss-of-function mutants *rh6-1*, *rh8-1* and *rh12-2* have reduced rosette size and fresh weight relative to the wild-type (the *SOV* deficient Col-0 background) (Fig. 1b,c **and** Supplementary Fig. 3). To test if they are functionally redundant, double *rh68* (*i.e.*, *rh6-1 rh8-1*), *rh612* and *rh812* mutants, and a triple *rh6812* mutant were generated. We found rosette growth is proportional to the copy number of functional *RH* genes, with the triple mutant *rh6812* displaying the most extreme diminished rosette diameter, fresh weight, petiole and leaf expansion, and a failure to flower (Fig. 1b,c). Independent hypomorphic *amiRH6 rh812* lines with strong partial suppression of *RH6* transcripts displayed less evident seedling alterations but had significantly reduced rosette growth (Supplementary Fig. 4e-h). These phenotypes are due to RH deficiency, as growth and reproduction of *rh6812* was rescued by a 3.3-kb genomic fragment of *RH6* with a C-terminal FLAG epitope tag (*gRH6-FLAG*) (Fig. 1b,c).

**Fig. 1.**
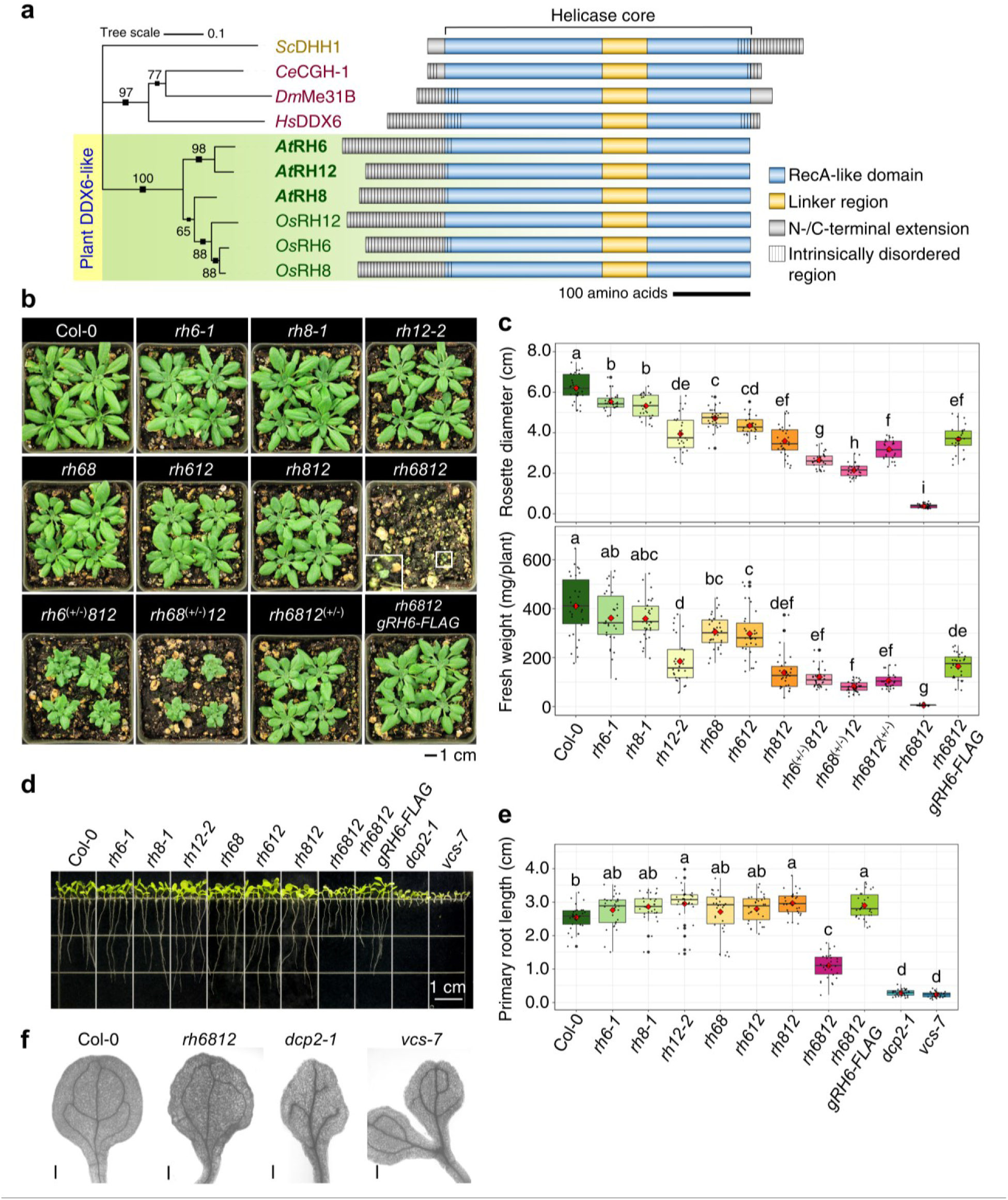
Arabidopsis *RH6*, *RH8* and *RH12* are functionally redundant and required for growth and development. **a**, Phylogenetic relationship and schematic of DHH1/DDX6-like proteins from the yeast *Saccharomyces cerevisiae* (*Sc*DHH1), roundworm (*Ce*CGH-1), fruit fly (*Dm*Me31B), human (*Hs*DDX6), *Arabidopsis* (*At*RH6*, At*RH8 *and At*RH12), and rice (*Os*RH6*, Os*RH8 *and Os*RH12). The tree is to scale, with branch lengths measured as the number of substitutions per site. Numbers on branches indicate bootstrap values. **b**, Rosette growth phenotype of 39-day-old plants of Col-0 wild-type in comparison to the single (*rh6-1*, *rh8-1* and *rh12-2*, double (*rh68*, *rh62* and *rh812*), double homozygous heterozygous (*rh6*^(+/-)^*812*, *rh68*^(+/-)^*12* and *rh6812* ^(+/-)^) and triple (*rh6812*; inset) mutant combinations, and a transgenic line homozygous for the *rh6812* triple mutant alleles with an introduced genomic *RH6* wild-type allele C-terminally tagged with the FLAG epitope (*rh6812 gRH6-FLAG*). Plants grown directly on soil; representative plants were selected. **c**, Rosette diameters (*n*=28) and fresh weight (*n*=30) of 39-day-old plants in **b**. **d**, 7-day-old seedlings. **e**, Primary root length (*n*=15) of seedlings in **d**. **f**, Representative images of the cotyledon vasculature of 7-day-old seedlings of Col-0 wild-type, *rh6812*, *dcp2-1* and *vcs-7* mutants. Bar = 0.3 mm. For **c** and **e**, means significantly different between genotypes are indicated by different letters (*p* < 0.05, ANOVA with Tukey’s HSD test).

Arabidopsis decapping mutants display seedling phenotypes including shortened primary roots^12, 14, 15^. The *rh* single and double mutants as well as *rh6812 gRH6-FLAG* homozygotes had seedling roots of similar or longer length than Col-0, whereas *rh6812* roots were significantly shorter (Fig. 1d,e **and** Supplementary Fig. 4b,c). However, *rh6812* roots were significantly longer than those of the decapping mutants *dcp2-1* and *vcs-7* (Fig. 1d,e). These three genotypes share chlorotic and small cotyledons with disorganized veins (Fig. 1f). We conclude that *RH6*, *RH8*, and *RH12* have additive redundant functions in regulating multiple aspects of growth and development.

Severe growth defects exhibited by mRNA decay mutants can be partly due to accumulation of siRNA produced from aberrant RNAs^17, 37^. Developmental defects of *dcp2-1* and *vcs-6* are partially rescued by mutation of *RNA-DEPENDENT RNA POLYMERASE 6* (*RDR6*)^17^. The *rh6812* growth phenotype is not primarily due to RDR6-mediated gene silencing, as neither the quadruple *rh6812 rdr6^sgs2–1^* nor the *rh6812 sgs3-1* (*suppressor of gene silencing 3-1*) genotype was rescued (Supplementary Fig. 5a,b).

### RH6, RH8 and RH12 are nucleocytoplasmic and associate with PBs and SGs

The subcellular localization of the three RHs was determined with transgenic lines that individually express genomic *RH6*, *RH8*, or *RH12* translationally fused with a C-terminal red fluorescent protein (*gRH6-RFP*, *gRH8-RFP*, or *gRH12-RFP*). All three RH-RFPs were nucleocytoplasmic in the root meristem region of seedlings grown under standard conditions (Fig. 2a,b **and** Supplementary Fig. 6a-f). The RH N-terminal region contains glutamine- and proline-rich sequences predicted to be intrinsically disordered (Fig. 1a), a property that facilitates formation of macromolecular condensates through liquid-liquid phase separation^38^. This was evaluated in the cortex cells of the root meristematic zone. Seedlings were bathed in water, placed under a coverslip and imaged within 2 min and again after 30 min. RH6-RFP was initially cytoplasmically dispersed and then present in large foci (Fig. 2c). These foci represent mRNP complexes as the pre-treatment of seedlings with cycloheximide inhibited the formation of RH6 foci altogether. Translation termination-dependent foci were similarly observed for RH8 and RH12 (Supplementary Fig. 6g-h). We conclude these RHs form mRNP assemblies with mRNAs released from polysomes.

**Fig. 2.**
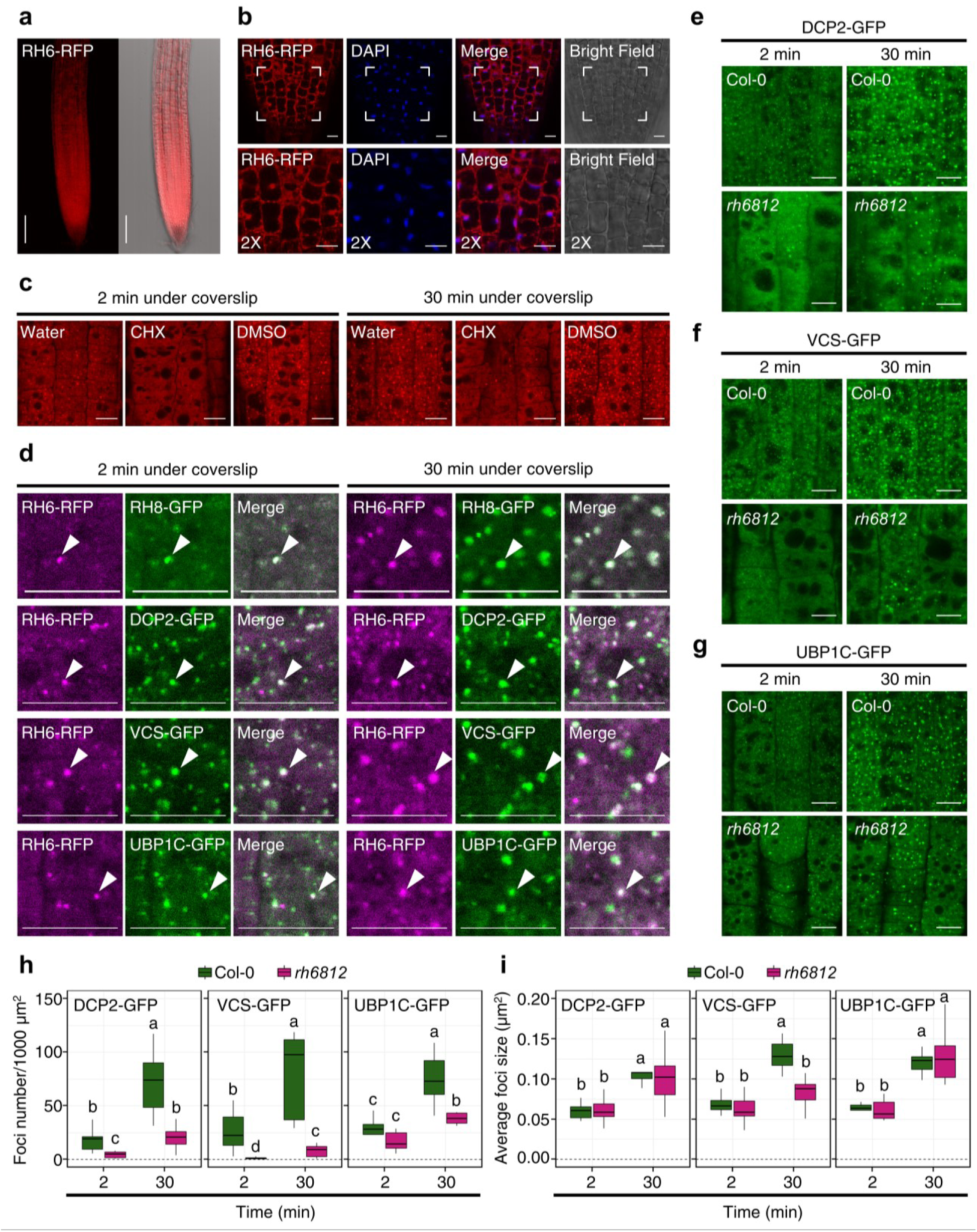
RH6, RH8 and RH12 form cytoplasmic mRNPs contributing to PB and SG formation. **a**, RFP fluorescence in root tissues of 4-day-old *genomic(g)RH6-RFP* seedlings. Left panel, maximum projection of 12 *z*-planes evenly spread across 55 µm; right panel, overlay on bright field image. Bars = 100 µm. **b**, Root meristem of seedlings in **a** counterstained with 4’,6-diamidino-2-phenylindole (DAPI) for nuclear visualization. Lower panels, magnification of framed areas of upper panels. **c**, Subcellular localization of RH6-RFP in the root meristem of 4-day-old seedlings submerged in water under a coverslip for 2 min. The same region re-imaged after 30 min. Seedlings pre-treated with 0.001% (v/v) dimethyl sulfoxide (DMSO) with or without cycloheximide (CHX) for 3 min by vacuum infiltration before imaging. Images representative of *n* ≥ 3 seedlings. **d**, Colocalization of RFP and GFP signals in the root meristem of 4-day-old seedlings co-expressing *gRH6-RFP* with *gRH8-GFP*, *proDCP2:DCP2-GFP*, *proVCS:VCS-GFP*, or *pro35S:UBP1C-GFP*. Imaging as described for **c**. RFP signals false-colored magenta. Arrowheads denote examples of foci with colocalized RFP and GFP signals. **e-g**, GFP fluorescence of DCP2-GFP, VCS-GFP, and UBP1C-GFP in the root meristem of Col-0 or *rh6812*. Developmental age-matched 4-day-old seedlings were imaged as in **c**. Images representative of *n* ≥ 6 seedlings per genotype. Bars in **b-g** = 10 µm. **h-i**, quantification of number and size of DCP2-GFP (*n*=9-12), VCS-GFP (*n*=10), and UBP1C-GFP (*n*=8) foci. Means significantly different between genotypes indicated by different letters (*p* < 0.05, ANOVA with Tukey’s HSD test).

Next, we considered if RH6, RH8, and RH12 are targeted to the same mRNPs. Root cortical cells coexpressing *gRH6-RFP* and *gRH8-GFP* colocalized in nearly all detectable cytoplasmic foci (Fig. 2d). The systematic evaluation of all three RHs confirmed all co-localize in the foci (Supplementary Fig. 7a-c). We asked whether RH-containing foci are PBs or SGs. By use of two GFP-tagged PB (DCP2 and VCS) markers, we resolved DCP2 assemblies after 2 min which enlarged after 30 min and often colocalized with RH6 (Fig. 2d **and** Supplementary Fig. 8a). The lack of complete overlap of the RH6 and DCP2 foci suggests heterogeneity in mRNP formation and dynamics. Similar colocalization dynamics were observed for RH8 and RH12 with DCP2 (Supplementary Fig. 8b,c) and VCS-GFP expressed under its native promoter (Fig. 2d **and** Supplementary Fig. 9a-c).

The incomplete association of the RHs with DCP2 and VCS led us to test whether the RHs overlap with the SG marker OLIGOURIDYLATE-BINDING PROTEIN 1C (UBP1C), a mammalian TIA-1/TIAR ortholog^6^. We confirmed overlap of UBP1C and RH foci within 2 min and increased after 30 min, in both foci size and number, in the root meristem of seedlings coexpressing the *gRH-RFPs* and *pro35S:UBP1C-GFP* (Fig. 2d **and** Supplementary Fig. 10a-c). Borders of the complexes were enriched for one or the other protein. This demonstrates that assemblies containing the three RHs dynamically associate with PBs as well as with SGs to form hybrid and heterogenous mRNP complexes.

### RH6, RH8 and RH12 contribute to the assembly of PB and SG complexes

To determine if RH function influences the formation of PB and SG assemblies, foci were examined in *rh6812* mutants expressing DCP2-GFP, VCS-GFP, or UBP1C-GFP. In contrast to DCP2 and VCS assembly dynamics in the root meristem of Col-0, these markers were in significantly fewer foci both at 2 and 30 min in *rh6812* (Fig. 2e,f,h). The size of DCP2 foci increased in both Col-0 and *rh6812* but VCS foci only increased in Col-0 (Fig. 2i). The deficiency in RHs also limited the temporal increase in number but not the size of UBP1C-containing SGs. These data provide further evidence of PB heterogeneity and indicate RHs contribute to the formation of large PBs and SGs.

### *RH6*, *RH8*, and *RH12* are required for transcriptome and translatome homeostasis

As the *rh6812* genotype had dampened PB and SG formation, we hypothesized that the RHs may contribute to regulation of transcript abundance and translation. We established the genotype *rh6812* expressing an epitope tagged RIBOSOMAL PROTEIN L18 that enables Translating Ribosome Affinity Purification (TRAP)^39^, to magnetically capture transcripts undergoing translation. This allowed comparison of *rh6812* and Col-0 seedling poly(A)^+^ mRNA (Total mRNA; transcriptome) and ribosome-associated poly(A)^+^ mRNA (TRAP mRNA; translatome) (Fig. 3a). RNA sequencing was performed on triplicate biological samples to an average depth of 19.4 million reads, revealing reproducible differences between the two genotypes and two mRNA populations (Fig. 3b, Supplementary Fig. 11a **and Supplementary Table 1**). Alterations in the transcriptome of the *rh6812* mutant was highly similar to alterations in its translatome (Pearson correlation 0.94) (Fig. 3c), with 409 and 419 mRNAs uniquely enriched and 151 and 666 mRNAs uniquely depleted in the Total and TRAP comparisons (*rh6812* relative to Col-0), respectively (Fig. 3d). The *rh6812* transcriptome and translatome were enriched in mRNAs with Gene Ontology (GO) functions associated with stress and innate immunity and depleted of mRNAs associated with general growth and primary metabolism, including polysaccharide metabolic process, rhythmic process, multidimensional cell growth, and regulation of hormone levels (Supplementary Fig. 11b **and Supplementary Table 2**).

**Fig. 3.**
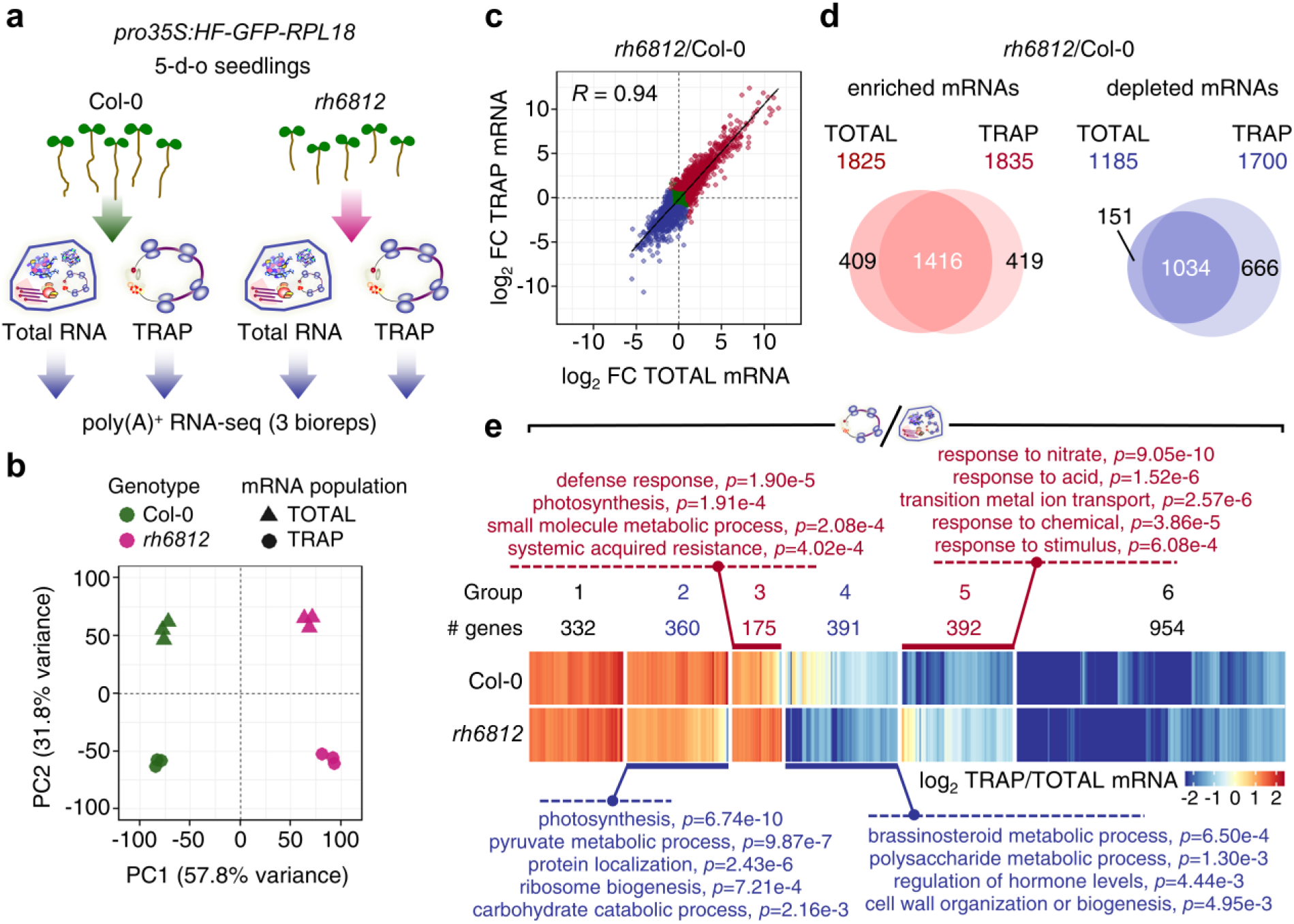
Attenuation of *RH6*, *RH8* and *RH12* function shifts the seedling steady-state transcriptome and translatome from a general growth to stress-responsive state. **a**, Schematic representation of experimental setup for transcriptome and translatome analyses. Ribosome-associated RNAs were isolated from developmental age-matched 5-day-old seedlings of Col-0 and *rh6812*, both expressing His-FLAG-GFP tagged RIBOSOMAL PROTEIN L18 (*pro35S:HF-GFP-RPL18*) using Translating Ribosome Affinity Purification (TRAP). PolyA+ RNA (Total) was isolated from the same tissue. **b**, Principal component (PC) analysis of edgeR-normalized read counts (CPMs) of Total and TRAP mRNAs from Col-0 and *rh6812.* c, Scatter plot comparison of the log_2_ FC between Total and TRAP mRNAs in *rh6812* relative to Col-0. *R* represents Pearson correlation coefficient of significantly enriched and depleted genes (red and blue dots, |log_2_ FC| ≥ 1, FDR < 0.01 in either RNA populations). **d**, Overlap of genes significantly enriched or depleted in *rh6812* relative to Col-0 (|log_2_ FC| ≥ 1, FDR < 0.01). **e**, Heatmap comparing 2604 genes with efficient or poor translation status in Col-0 or *rh6812* (|log_2_ TRAP/Total mRNA| ≥ 1, FDR < 0.01). These were classified into 6 groups based on the patterns of their translational status between the two genotypes. Number of genes in each group is shown. Representative GO terms enriched for each group sets and their adjusted *p*-values given.

Next, we investigated the impact of RH deficiency on translational status, the proportion of individual polyA^+^ mRNAs associated with ribosomes. Translation status of mRNAs varied in both Col-0 and *rh6812* seedlings (Supplementary Fig. 11c **and Supplementary Table 1**). Focusing on the 2,604 mRNAs with optimal and suboptimal translation in at least one genotype (|log_2_ FC| ≥ 1, FDR < 0.01), we resolved groups with similar (groups 1 and 6) or distinct (groups 2-5) translational status in the two genotypes (Fig. 3e **and Supplementary Table 3**). mRNAs with higher translation status in Col-0 than in *rh6812* were associated with primary metabolism, protein localization and ribosome biogenesis in group 2 and brassinosteroid metabolic process, polysaccharide metabolic process, and cell wall organization or biogenesis in group 4. Conversely, higher translational status was evident in *rh6812* for mRNAs associated with the defense response and systemic acquired resistance in group 3, and responses to different stimuli in group 5. This demonstrates the RHs contribute to transcriptome and translatome homeostasis under non-stress conditions by directly or indirectly promoting translation of mRNAs associated with growth and development and limiting ectopic translation of mRNAs involved in cellular response to stress and external stimuli.

### RH6, RH8, and RH12 facilitate decay of select short-lived mRNAs

The colocalization of the RHs with the decapping proteins DCP2 and VCS prompted an evaluation of their role in decay of stress/stimuli-responsive and other mRNAs. The experiment commenced with infiltration of *rh6812* and Col-0 seedlings with the transcription inhibitor cordycepin, sampling after 0, 15, 30, 60 and 120 min, and RNA-sequencing of rRNA-depleted total RNA (Fig. 4a). Triplicate biological samples were sequenced to an average depth of 22.7 million reads. After data normalization and exclusion of low abundance mRNAs, transcript abundances were used to model decay rates for each gene and genotype^11^.

**Fig. 4.**
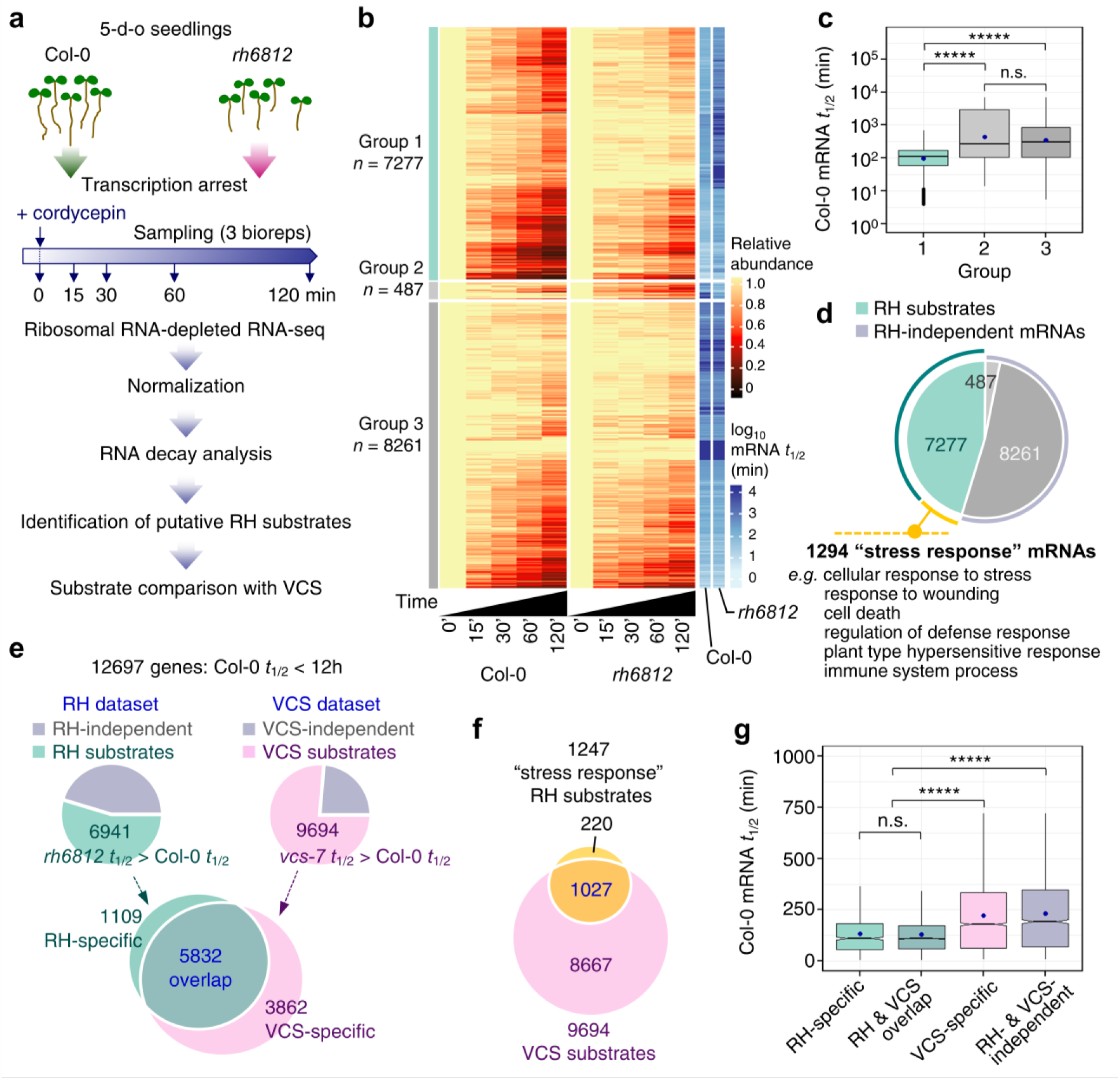
RHs facilitate decay of short-lived mRNAs. **a**, Experimental design for global RNA decay analysis. Seedlings were sampled at 5 time after cordycepin treatment in triplicate. RNA-seq libraries were generated from total RNA following ribosomal RNA depletion. **b**, Heatmap of mRNA decay over 120 min and *t*_1/2_ of 16,025 transcripts in Col-0 and *rh6812*. mRNA decay is presented as change in mean relative mRNA abundance and hierarchically clustered into three groups. Group 1 mRNAs decreased decay rates in *rh6812* (putative RH substrates, *rh6812 t*_1/2_ > Col-0 *t*_1/2_); group 2 mRNAs increased decay rates in *rh6812* (*rh6812 t*_1/2_ < Col-0 *t*_1/2_); group 3 mRNAs unaffected (*rh6812 t*_1/2_ ≃ Col-0 *t*_1/2_). **c**, Comparison of Col-0 mRNA *t*_1/2_ between each group in **b**. **d**, Pie chart highlighting proportion of RH substrates identified in **b** involved in stress or defense responses based on GO enrichment. **e**, Pie charts of proportion of putative substrates of RH (group 1) and VCS; Venn diagrams showing the overlap between the populations. RH and VCS substrates are stabilized in *rh6812* and *vcs-7*, respectively. *vcs-7* mRNA decay data were modeled exactly as for RH, with comparison limited to genes with Col-0 mRNA *t*_1/2_ < 12 h. **g**, Venn diagram comparing overlap of RH substrates related to “stress response” with VCS-dependent decapping pathway substrates. **h**, Plots comparing the Col-0 *t*_1/2_ of RH-substrates, RH and VCS substrates classified in **f**. In **c** and **g**, ***** *p* < 2.2e-16; n.s., not significant (Wilcoxon test).

We found that loss of the RHs increased stability of otherwise rapidly degraded transcripts, with the minimum and median mRNA half-life (*t*_1/2_) increased from 3.8 and 160.9 min in Col-0 to 5.2 and 314.1 min in *rh6812*, respectively (Supplementary Fig. 12). This was also evident when the mRNAs were classified into three groups by relative effect of the RH deficiency on *t*_1/2_ (Fig. 4b **and Supplementary Table 4**). Group 1 (*n* = 7,277) mRNAs decayed more slowly in *rh6812* (*rh6812 t*_1/2_ > Col-0 *t*_1/2_). These are termed RH substrates, as their decay is directly or indirectly dependent on the presence of RHs in Col-0 for their rapid turnover. Group 2 (*n* = 487) mRNAs decayed more rapidly in the triple mutant (*rh6812 t*_1/2_ < Col-0 *t*_1/2_) and group 3 mRNAs were unaffected (*n* = 8,261; *rh6812 t*_1/2_ ≃ Col-0 *t*_1/2_).

The RH substrates (group 1) had characteristics that provide insights into RH function and biological relevance. First, these collectively had the shortest half-lives in Col-0 of the three groups (Fig. 4c). Their median *t*_1/2_ in Col-0 was 108 min, whereas that of groups 2 and 3 were 263 and 299 min, respectively. Second, the RH substrates had fewer codons, typically fewer introns and were shorter than non-substrate mRNAs, whereas their 5’ and 3’ untranslated region lengths showed no overall difference (Supplementary Fig. 13a,b). Third, the RH substrate GO functions encompassed a wide range of biological processes (**Supplementary Table 5**). Of these,1,294 (17.8%) are involved in stress and immune responses (Fig. 4d). These “stress response” mRNAs include disease resistance proteins, transcription factors, and signaling proteins such as kinases (**Supplementary Table 6**).

Since all three RHs colocalize with DCP2 and VCS, we hypothesized that their substrates are degraded in a decapping-dependent manner. A parallel decay experiment was performed in the decapping mutant *vcs-7*^11^, and co-analysis of *rh6812* and *vcs-7* effects on decay identified 5,832 out of 6,941 RH substrates (84%) as VCS substrates (Fig. 4e). The analysis also identified RH-specific and VCS-specific substrates. Comparison of 1,247 “stress response” RH substrates with those dependent on VCS revealed that the majority (1,027, 82.4%) require VCS and are therefore degraded by the decapping pathway (Fig. 4f). The RH substrates had a shorter *t*_1/2_, whether or not they require VCS, as compared to those that only require VCS or neither RHs nor VCS for decay (Fig. 4g). These results verify that the majority of mRNAs targeted by the RHs are ephemeral targets of 5’-to-3’ turnover and provide further evidence that the RHs accelerate the decay of a subset of mRNAs.

### Stress-related mRNAs are stabilized and effectively translated in the *rh6812* genotype

We also considered if steady-state abundance and translation of RH substrates was altered in *rh6812* seedlings relative to Col-0. Meta-analysis of the combined mRNA decay, transcriptome and translatome datasets revealed that of all mRNAs with higher levels in *rh6812* Total and TRAP mRNA populations, 42% (383) were RH substrates (Fig. 5a). A homodirectional increase in steady-state and translated mRNA abundance was expected, but the 383 RH substrates also had a significantly higher translational status than the 522 RH-independent mRNAs from this group (Fig. 5b). This indicates that increased stability and abundance of these RH substrates is coupled with their preferential ribosome association in the *rh6812* genotype, which was not evident for the full cohort of RH substrates. GO enrichment analysis revealed that the 383-mRNA cohort is enriched for transcripts encoding stress/defense response proteins (**Supplementary Table 7**). The decay kinetics for eight representative defense genes (Fig. 5c) support the conclusion that these RH substrates are synthesized at a basal level but rapidly degraded under standard growth conditions.

**Fig. 5.**
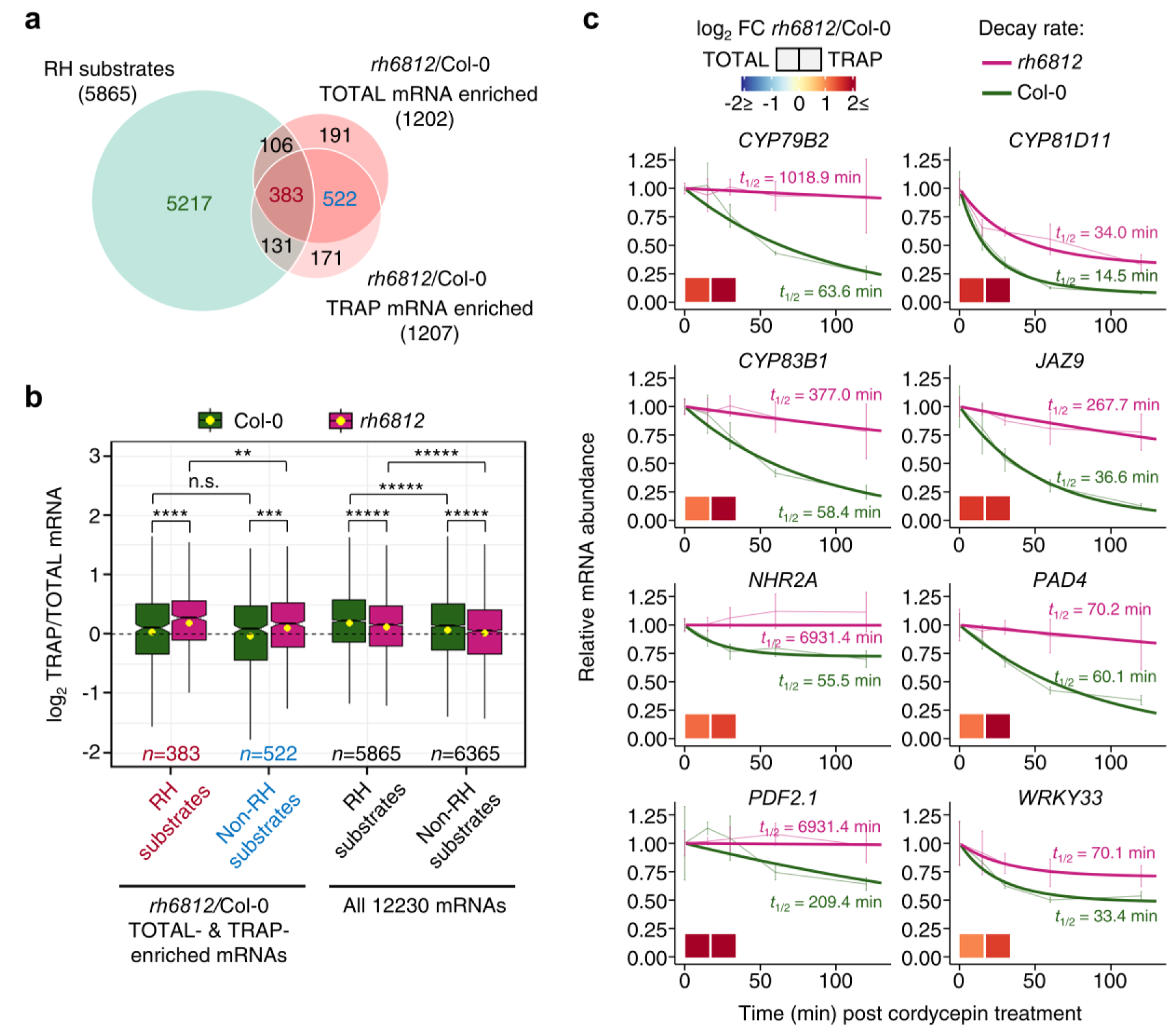
Stabilization of stress response mRNAs in the *rh6812* genotype is associated with an increase in the abundance and translational status. **a**, Venn diagram showing the overlap between RH decay substrates and steady-state TOTAL and TRAP mRNAs that are enriched in *rh6812* relative to Col-0 (log_2_ FC ≥1, FDR < 0.01) from a combined analysis of mRNA decay, transcriptome and translatome data comprised of 12230 genes. **b**, Box plot comparing translational status (log_2_ TRAP/TOTAL mRNA) of RH substrates and RH-independent mRNAs. ***** *p* < 2.2e-16; **** *p* = 1.2e-13; *** *p* = 3.4e-9; ** *p* = 0.011; n.s., no significant difference (Wilcoxon test). **c**, Decay profiles of representative stress and defense response mRNAs with an increase in TOTAL and TRAP mRNA abundance (log_2_ FC indicated by colored boxes) and translational status in *rh6812* relative to Col-0. mRNA decay profile of each genotype is presented as relative mRNA abundance after inhibition of transcription by cordycepin treatment (thin line, mean ± SE, *n*=3) along with modelled value (thick line). *t*_1/2_ for each genotype is shown.

### The *rh6812* mutant phenotype is associated with autoimmunity

The observation that aseptically grown *rh6812* seedlings stabilize and actively translate mRNAs involved in defense response led us to hypothesize that the diminutive mutant phenotype results from a constitutive immune response. The most ectopically elevated mRNAs in *rh6812* included many well-known defense response genes (Fig. 6a). Moreover, levels of the defense phytohormone salicylic acid (SA) were significantly elevated in *rh6812* relative to Col-0 seedlings (Fig. 6b), consistent with increased levels of *ISOCHORISMATE SYNTHASE 1* (*ICS1*) mRNA, encoding a key enzyme in SA biosynthesis (Fig. 6a). Transcript levels of the defense response marker *PATHOGENESIS-RELATED 1* (*PR1*) were also significantly elevated in *amiRH6 rh812* lines, but to lower levels than in *rh6812* seedlings (Fig. 6c). Both *amiRH6 rh812* and *rh6812* seedlings were more resistant to the pathogenic Noco2 isolate of the oomycete *Hyaloperonospora arabidopsidis* than Col-0, with the *rh6812* genotype showing the highest level of resistance (Fig. 6d,e).

**Fig. 6.**
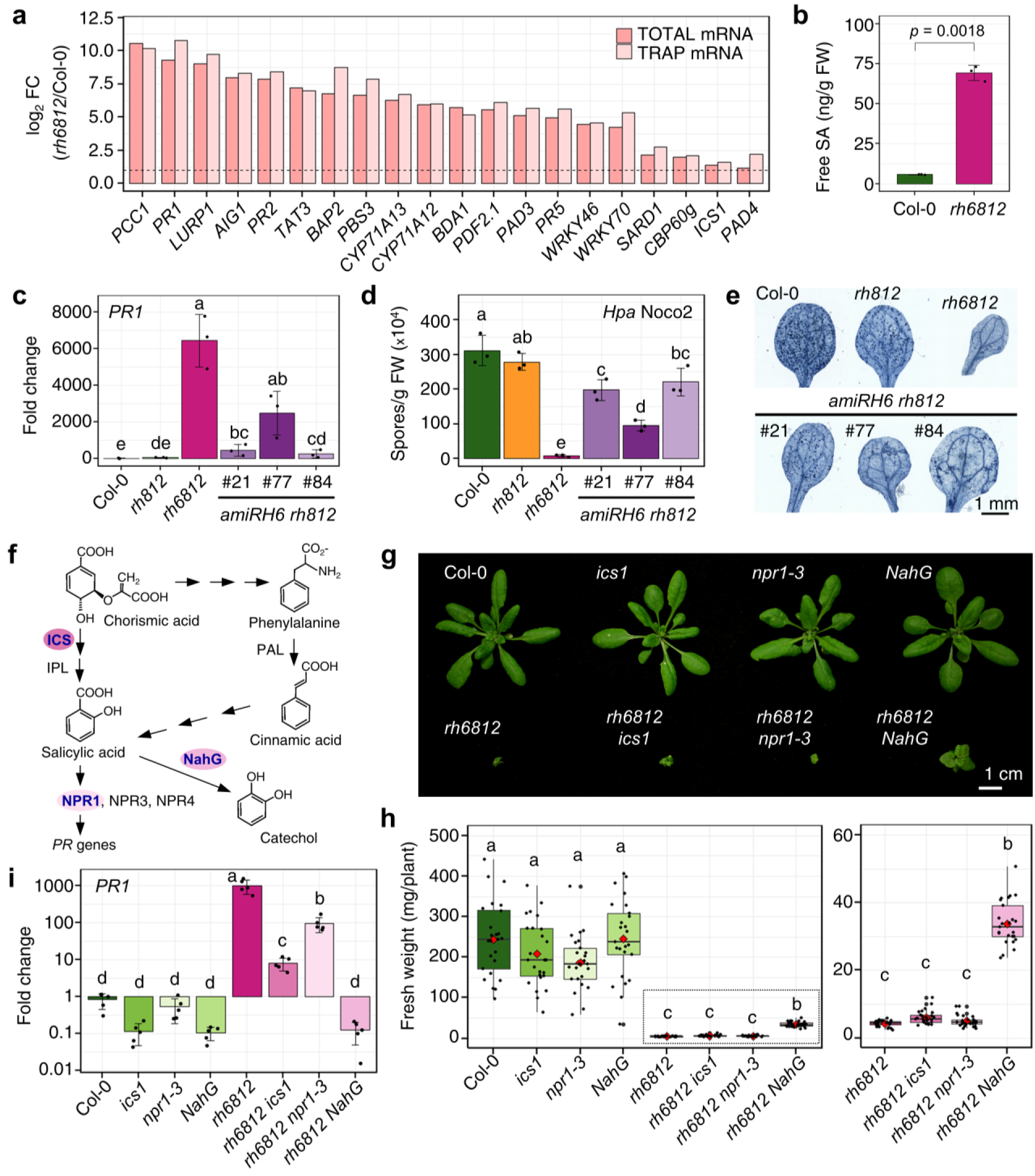
The triple *rh6812* mutant exhibits a constitutive immune response. **a**, Examples of defense-related genes with constitutively elevated transcripts in *rh6812*. Data are presented as log_2_ FC of Total and TRAP mRNA abundance in *rh6812* relative to Col-0. Horizontal dashed line, log_2_ FC = 1. **b**, SA content in 7-day-old seedlings of *rh6812* in comparison to Col-0. Error bars, SD (*n*=3); Welch Two Sample t-test. **c**, RT-qPCR quantitation of *PATHOGENESIS-RELATED GENE 1* (*PR1*) transcripts in 7-day-old seedlings (*n*=3) of Col-0, *rh812*, *rh6812* and *amiRH6 rh812* lines. Fold-change calculated using *Actin 1* (*ACT1*) as reference. Error bars, SD (*n*=3). Experiment repeated twice with similar results. **d**, Quantification of *Hpa* Noco2 sporulation on seedlings. **e**, Representative cotyledons stained with lactophenol trypan blue at 7 days after inoculation. **f**, Diagram of SA biosynthesis, perception and degradation. **g**, Representative rosette growth phenotype of 28-day-old plants: Col-0, single *ics1* and *npr1-3* mutants, Col-0 overexpressing the bacterial *NahG*, triple *rh6812* mutant, quadruple *rh6812 ics1* and *rh6812 npr1-3* mutants, and triple *rh6812* mutant combined with *NahG*. **h**, Fresh weight of 28-day-old plants (*n*=25) of genotypes presented in **g**. **i**, RT-qPCR quantitation of *PR1* transcript levels in 7-day-old seedlings (*n*=5) of genotypes presented in **g**. Relative transcript fold-change calculated using *ACT1* as a reference. Means significantly different between genotypes in **c**, **d**, **h** and **i** are indicated by different letters; **h** and **i** data were log transformed (*p* < 0.05, ANOVA with Tukey’s HSD test).

Autoimmunity was demonstrated previously for the mRNA decay mutants *up-frameshift 1* (*upf1*)^40^ and *pat1*^16^. Their autoimmune phenotypes can be uncoupled by inactivation of the immune regulator ENHANCED DISEASE SUSCEPTIBILITY 1 (EDS1) and/or PHYTOALEXIN DEFICIENT 4 (PAD4)^16, 23^. We found that *PAD4* transcripts were stabilized with concomitant elevation in Total and TRAP mRNA abundance as well as translation status in *rh6812* seedlings (Fig. 5d **and** Fig. 6a). This coincided with elevation of known EDS1/PAD4-inducible mRNAs in the transcriptome and translatome of *rh6812* (Supplementary Fig.14a). However, based on phenotypic analyses and quantification of *PR1* transcripts and SA levels in the quadruple mutants *rh6812 eds1-2* and *rh6812 pad4-1* (Supplementary Fig.14b-e), the constitutive immunity exhibited by *rh6812* is partially dependent on PAD4 but not EDS1 function. We tested further whether manipulation of SA synthesis, perception, or catabolism would rescue the *rh6812* phenotypes (Fig. 6f). This included combining *rh6812* with mutations of *ICS1* and *NONEXPRESSER OF PR GENES 1* (*NPR1*) and ectopic expression of the bacterial salicylate hydroxylase *NahG*. Although the homozygous *rh6812 ics1* and *rh6812 npr1-3* genotypes were as diminutive as the triple mutant, *NahG* partially rescued *rh6812* rosette growth (Fig. 6g,h). This rescue was accompanied by a significant reduction in *PR1* transcripts in *rh6812 NahG* seedlings, to levels indistinguishable from the four control genotypes Col-0, *ics1*, *npr1-3* and *NahG* (Fig. 6i). *PR1* transcript levels were reduced in *rh6812 ics1* and *rh6812 npr1-3* relative to *rh6812* seedlings, but remained significantly higher than in the control genotypes. We conclude that severe RH deficiency promotes a SA-mediated immune response that contributes to dampening of growth in *rh6812*. Collectively, these data uncover a functional role of the RHs in limiting basal accumulation of transcripts associated with innate immunity.

## Discussion

The Arabidopsis DHH1/DDX6-like proteins RH6, RH8, and RH12 are integral for growth and development, with the triple *rh6812* mutant showing pleiotropic phenotypes with some overlap with other mRNA decapping mutants. When expressed under the control of their native promoters all three RHs localize to the nucleus and cytoplasm, consistent with prior analyses using ectopic expression^41, 42^. SGs and PBs are the two major conserved translationally repressed mRNP complexes in the cytoplasm of eukaryotic cells, with PBs typically associated with translational repression and 5’-decapping dependent decay^43, 44^. In the root meristematic region of seedlings, the RHs form inducible mRNP complexes that overlap with one another and with the PB markers DCP2 and VCS and the SG marker UBP1C. Consistent with this finding, RH6 interacts with the translational repressor and decapping activator DCP5, while all three RHs co-immunoprecipitate with the nonsense-mediated decay factor UPF1, along with the core decapping activators DCP1 and VCS, the eukaryotic translation initiation factor eIF4G, and several poly(A)-binding proteins^42^.

We found that the RHs and PB and SG markers are not uniformly overlapping, strengthening the notion that these complexes are heterogeneous. Our demonstration that RHs are required for the formation of visible DCP2 and VCS foci suggests that these helicases contribute to PB assembly and dynamics as demonstrated for DHH1/DDX6 proteins in other organisms^31, 32, 45, 46^. It remains elusive as to how RHs contribute to the formation of DCP2 and VCS assemblies, while only a subset of them overlap with the RHs. In addition to PBs, the RHs may contribute to the formation of stress-induced UBP1C SGs, as observed in yeast DHH1^38^, although this might be lineage-specific because depletion of DDX6 in human HeLa cells does not affect SG formation^45^.

In Arabidopsis, depletion of PBs in the *dcp5-1* mutant results in precocious translation of specific mRNAs in etiolated seedlings^47^. Here we found that reduction of PBs in light-grown *rh6812* seedlings is associated with ectopic translation of stress- and defense-response mRNAs. This suggests that the role of PBs in translational repression may include the control of stress-related mRNAs under non-stress conditions. Furthermore, our global RNA decay analysis showed that turnover of thousands of short-lived mRNAs is compromised in *rh6812* seedlings. The co-localization and 84% overlap of RH and VCS substrates strongly supports the conclusion that the RHs are components of the decapping-dependent, 5’-to-3’ decay machinery. Comparison of RH and VCS substrates also supports the notion that mRNA decay does not necessarily occur in visible PBs, as their formation was impaired in *rh6812* but degradation of over 3,800 VCS-specific substrates was unaffected. It is yet to be determined whether RH-assisted decay occurs in PBs of smaller than detectable size, or cotranslationally on elongating ribosomes.

Plants have evolved diverse and intricate systems to rapidly sense and respond to environmental insults including pathogens. While lacking adaptive immunity, plants have a battalion of immune receptors including resistance (*R*) genes that encode nucleotide-binding leucine-rich repeat (NLR) proteins to recognize different pathogen-derived effector proteins and trigger a robust defense response^48, 49^. Since defense occurs at the expense of cellular energy reserves, immune response gene and protein activation must be tightly regulated to avoid autoimmunity and maintain plant growth. This includes NLR regulation that operates from epigenetic to post-translational levels^48, 50^. Post-transcriptional control of *R* gene transcripts is multi-faceted, including regulation of alternative splicing^51, 52^, alternative polyadenylation^53^, small RNA-mediated silencing^54^, nonsense-mediated decay^23^, and translation^55^. The dysregulation of any of these processes often causes a sensitized or constitutive immune response.

There is compelling evidence that general cytoplasmic mRNA decay contributes to the control of the defense response in Arabidopsis. Pattern-triggered immunity activates decapping-dependent turnover of transcripts known to decline in response to the elicitor flg22^56^, although the effect on global decay is unknown. Moreover, the decapping mutant *pat1* exhibits an autoimmune phenotype that is suppressed by mutations of the NLR SUPPRESSOR OF MKK1 MKK2 2 (SUMM2) and the defense regulator EDS1^16^. In yeast, PAT1 interacts with DHH1 to repress translation and activate mRNA decapping^35, 57^. Similar to *pat1*, the triple *rh6812* mutant exhibits autoimmunity albeit in an EDS1-independent manner. Our mRNA decay analysis identified 44 annotated *R* gene transcripts along with over a thousand other stress- and defense-response mRNAs, including those encoding plant-specific transcription factors that are stabilized in the *rh6812* genotype. Many of these mRNAs increase in translational status in consort with elevated stability and abundance. This indicates that RHs actively repress basal or aberrant activation of the plant immune response. We hypothesize that the DHH1/DDX6-like protein family plays a conserved role as a negative regulator of innate immunity in eukaryotes, as the depletion of human DDX6 induces global upregulation of interferon-stimulated and other immune genes in the absence of infection^58^. The deficiency in RH activity is accompanied with a reduction in plant growth, reminiscent of genotypes with constitutive innate immunity^59^. While the *rh6812* phenotype is partially rescued when ectopic accumulation of the key defense metabolite SA is controlled, we cannot exclude the role of stabilized mRNAs unrelated to SA production in the reduction of rosette growth, given that RH substrates encompass diverse stress-response mRNAs and RH deficiency shifts the balance of stress and growth-related transcripts.

In summary, a severe deficiency of RH function results in catastrophic alteration of seedling mRNA homeostasis, allowing the accumulation and translation of stress- and defense-response transcripts, while dampening mRNAs essential for growth and fecundity. The regulation of mRNA turnover is integral to rapid changes in gene expression in response to stress^60^. This study illuminates the significant contribution of mRNA decay in shaping the cellular transcriptome and translatome under optimal conditions. The findings raise the notion that stress-responsive transcripts are constitutively synthesized at a basal level, but their accumulation and translation are limited by degradation. Inhibition of the decay machinery, in conjunction with activation of their transcription above the basal level, would allow a rapid and robust response to stress. This posits new questions as to whether RH function is dynamically modulated to allow stress-response transcripts to escape destabilization in response to cues that activate their transcription to provide pathogen resistance or stress resilience.

## Methods

Methods, including statements of data availability and any associated accession codes and references, are available in the **Supplemental Materials**.

## Acknowledgements

We thank Dr. Jane Parker for kindly providing *eds1-2* and *pad4-1* seeds, Dr. Martin Crespi for *rdr6^sgs2–1^* and *sgs3-1*, and Dr. Yuichiro Watanabe for *proDCP2:DCP2-GFP* in *dcp2-1.* We thank members of the Bailey-Serres and Sieburth laboratories for thoughtful discussion. This research was supported by United States National Science Foundation grants no. MCB-1021969 to J.B.-S. and L.S. and MCB-1716913 to J.B.-S. and a UC MacArthur Foundation Chair award to J.B.-S.T.C. was supported by a Royal Thai Government’s Development and Promotion of Science and Technology Talents Project scholarship.

## Author contributions

T.C., R.S., L.S. and J.B.-S. conceived and designed experiments; T.C., R.S., H.K. and K.A. performed experiments; A.K. and D.C. contributed to the preparation of genetic material; K.D., T.E., L.S. and J.B.-S. provided reagents; T.C., R.S., M.H., L.S. and J.B.-S. analyzed data. T.C., R.S., L.S. and J.B.-S. wrote the manuscript.

## Competing Interests

The authors declare no competing interests.

## Supplementary Materials

### Methods and Figures

**Supplementary Table 1:** Differential expression analysis of 14,391 genes (rpkm >5) by edgeR of TOTAL and TRAP mRNA-seq data associated with Fig. 3 and Supplementary Fig. 11.

**Supplementary Table 2:** Enriched GO categories of Total and TRAP mRNA differentially expressed in the triple *rh6812* mutant relative to the wild-type Col-0.

**Supplementary Table 3:** Enriched GO categories associated with each mRNA group in Fig.3e.

**Supplementary Table 4:** Initial (alpha) decay rate, mRNA half-life (*t*_1/2_), decay group, statistical parameter, and relative mRNA abundance of 16,025 genes associated with Fig. 4

**Supplementary Table 5:** Enriched GO terms associated with 7277 RH substrates

**Supplementary Table 6:** List of “stress response” RH substrates

**Supplementary Table 7:** Enriched GO terms associated 383 RH substrates with increased Total and TRAP mRNA abundance in *rh6812*

**Supplementary Table 8:** Oligo DNA sequences used for PCR-based genotyping, RT-PCR, and RT-qPCR in this study

## Supplemental Materials

### Materials and Methods

#### Plant materials and growth conditions

*Arabidopsis thaliana* mutants and transgenics are in the Columbia-0 (Col-0) ecotype (*SOV* deficient). The T-DNA insertion mutants *rh6-1* (Sail_111_H08), *rh8-1* (Salk_016830C), and *rh12-2* (Salk_016921C), *dcp2-1*^1^ (Salk_000519), *vcs-7*^2^ (Salk_032031), and *ics1*^3^ (SALK_133146C) were obtained from the Arabidopsis Biological Resource Center. The homozygous double *rh68*, *rh612* and *rh812*, and triple *rh6812* mutant combination lines were generated by crossing of the *rh6-1*, *rh8-1* and *rh12-2* alleles. The mutants *rdr6^sgs2-1^*, *sgs3-1*, *eds1-2*, *pad4-1*, *npr1-3* were described previously^4–7^. The transgenics *NahG*, *proDCP2:DCP2-GFP* in *dcp2-1*, *proVCS:VCS-GFP*, *pro35S:UBP1C-GFP* in *ubp1c-1* and *pro35S:HF-GFP-RPL18* were described previously^8–12^. Genotyping primers used are listed in **Supplementary Table 8**.

Seeds were surface-sterilized by incubation in 70% (v/v) ethanol for 5 min, 20 % (v/v) household bleach plus 0.01 % (v/v) Tween-20 for 5 min, and rinsed 5 times with sterile water. Seeds were plated on sterile solid Murashige and Skoog (MS) medium containing 0.5x MS salts (Caisson Laboratories), 0.5 % (w/v) Sucrose, 0.4 % (w/v) Phytagel (Sigma-Aldrich) [pH 5.7-5.8], followed by stratification at 4°C in complete darkness for 2 days before vertically grown in a growth chamber under a 16-h light (∼100 μmol photons·s^-1^·m-^2^) and 8-h dark cycle at constant 23°C. 7-day-old seedlings were transferred to soil containing Sunshine LC1 soil mix (JM McConkey) with 1.87 g/L Marathon insecticide (Crop Production Services, Riverside, California) and 1.4 g/ L osmocote 14-14-14 fertilizer, and grown in a growth room under long-day conditions (∼100 μmol photons·s^-1^·m-^2^). Alternatively, seeds were directly sown in soil, and stratified at 4°C in the dark for 2 days before being transferred to a growth room and grown under the same conditions. Plants were imaged using a Canon camera (Canon USA). Images were processed using Fiji software^13^.

Sterilized seeds of *rh6*^(+/–)^*812 pro35S:HF-GFP-RPL18* genotype plated on sterile solid medium were transferred after stratification to the growth chamber 12 h in advance of the *pro35S:HF-GFP-RPL18* seeds to compensate for their delayed germination. To prevent disturbance of seedlings and hasten tissue collection, seedlings of wild-type phenotype were removed from the segregating population of *rh6*^(+/-)^*812 pro35S:HF-GFP-RPL18* progeny after 4 days. Five-day-old seedlings of the *pro35S:HF-GFP-RPL18* and *rh6812 pro35S:HF-GFP-RPL18* genotypes were harvested and flash frozen in liquid nitrogen, and stored at −80°C until used. All sample preparations were carried out in at least three independent biological replicates.

#### Database search and *in silico* sequence analysis

DHH1/DDX6-like proteins from the yeast (*Saccharomyces cerevisiae*), roundworm (*Caenorhabditis elegans*), fruit fly (*Drosophila melanogaster*), and human (*Homo sapiens*) were downloaded from *Saccharomyces* Genome Database, WormBase, FlyBase, and GeneCards Human Database, respectively. Putative DHH1 orthologs from fungi were identified using the orthoDB database^14^, and their corresponding amino acid sequences were downloaded from UniProt database. Amino acid sequences of DDX6 orthologs from vertebrates were downloaded from the Emsembl genome browser. Amino acid sequences of DHH1/DDX6-like proteins from green algae and land plants were obtained from the Phytozome database following a BLASTP search of the yeast DHH1 sequence. In total, full-length amino acid sequences of 58 representative DHH1/DDX6-like proteins from fungi (12), animals (14), green algae (5), and land plants (27) were aligned using the MUSCLE function in the MEGA7 software^15^. The evolutionary history was inferred by using the Maximum Likelihood method following the Le_Gascuel_2008 model with 1000 bootstrap replicates^16^. Initial tree(s) for the heuristic search were obtained by applying Neighbor-Join and BioNJ algorithms to a matrix of pairwise distances estimated using Jones-Taylor-Thorton (JTT) model^17^. A discrete Gamma distribution was used to model evolutionary rate differences among sites. All positions with less than 95% site coverage were eliminated, with a total of 403 positions in the final dataset. The tree was visualized and drawn using the Interactive Tree Of Life (iTOL) software^18^.

#### Total RNA isolation, RT-PCR and quantitative RT-PCR

Total RNA was extracted from indicated tissues using TRIzol^TM^ reagent (Invitrogen), and purified using the Direct-zol^TM^ RNA MiniPrep (Zymo Research) according to the manufacturer’s instructions. Total RNA was treated with DNase I (New England Biolabs), and cDNAs were prepared from 2 μg of DNase-free total RNA using an oligo(dT) primer or gene-specific/stem-loop primers (Integrated DNA Technologies) and Maxima Reverse Transcriptase (Thermo Fisher Scientific) following the manufacturer’s instructions. End-point PCR was performed using 3 µL of 5x diluted cDNA solution with Taq DNA polymerase (G-Biosciences) and the appropriate primers according to the manufacturer’s instructions. For RT-qPCR analysis, amplification was performed in technical duplicate in the CFX Connect™ Real-Time PCR Detection System (Bio-Rad Laboratories) using iQ™ SYBR^®^ Green Supermix (Bio-Rad Laboratories) in a 15-µL reaction volume with 0.67 µm of each primer and 2 µL of 5× diluted cDNA. Relative transcript fold-change was calculated by the ΔΔCt method^19^. Primers used in RT-PCR and RT-qPCR are listed in **Supplementary Table 8**.

#### Plasmid construction and complementation of the triple *rh6812* mutant

A 3.3 kb *RH6* genomic fragment, spanning the upstream intergenic (promoter, 836 bp upstream of the start codon) region and gene transcript sequence without the stop codon, was amplified from Col-0 wild-type genomic DNA using the primer pair proRH6-F (5’CAC CTT TCT CTC TTT CTT TCG GAT GTT A-3’) and RH6(-stop)-R (5’-CTG ACA GTA GAT TGC CTT GT-3’) and introduced into pENTR/D-TOPO (Invitrogen) to generate a genomic RH6 entry clone. The DNA fragment was then introduced into the gateway-FH-OCST (nptII) T-DNA vector ^11^ to generate C-terminally FLAG tagged RH6 under its native promoter (*gRH6-FLAG*) via LR reaction. The construct was transformed into *Agrobacterium tumefaciens* strain GV3101 and then transformed into *rh6812*^(+/-)^ plants by the floral dip method^20^. Transgenic T1 and T2 seeds were selected on 0.5x MS medium supplemented with 50 μg/mL kanamycin.

#### Generation of transgenic plants and crossing

An artificial miRNA targeting *RH6* transcripts was designed using the WMD3-Web MicroRNA Designer^21^. The 404-bp *MIR319a* precursor was engineered by replacing the original *miR319a* with the artificial sequence targeting *RH6* (amiRH6, 5’-TTA ATA TTG GGT AAC ACC CAG-3’), and the *miR319a\** with *amiRH6\** (5’-CTA GGT GTT ACC CTA TAT TAT-3’). The engineered *MIR319a-amiRH6* precursor with the *attB1* and *attB2* flanking sequences was synthesized and placed in the pIDTSMART-AMP vector (Integrated DNA Technologies). The *MIR319a-amiRH6* sequence was transferred by *in vitro* recombination into pDONR-zeo using BP clonase (Invitrogen), and then further transferred into the pMDC32 binary vector^22^ using LR clonase (Invitrogen). The resulting *pro35S:amiRH6* construct was introduced into the double *rh812* mutant via *A. tumefaciens* GV3101 using the floral dip method^20^. T1 and T2 seeds were selected on 0.5x MS medium supplemented with 0.5% (w/v) sucrose and 50 μg/mL Hygromycin B (Sigma).

A 5.0 kb fragment of *RH8* genomic region (including 2.2 kb upstream of the start codon and the gene body without the stop codon) was amplified from of Col-0 wild-type genomic DNA using the primer pair proRH8-F (5’-ggg gac aag ttt gta caa aaa agc agg ctC TCT ACG GCG ATT GAT CTA AGC-3’) and RH8(-stop)-R (5’-ggg gac cac ttt gta caa gaa agc tgg gtA TTG GCA ATA AAT TGC CTG ATC G-3’) and introduced into pDONR-zeo vector (Invitrogen) to generate a genomic *RH8* entry clone. Similarly, a 3.5 kb *RH12* genomic region spanning 992 bp upstream of the start codon and the genic region without the stop codon was amplified from of Col-0 wild-type genomic DNA using the primer pair proRH12-F (5’CAC CTC AAG GTT TGT TTT GCC ATC A-3’) and RH12(-stop)-R (5’-ATC GAT CAA GCA ATC TAC TGT CAG-3’) and introduced into pENTR/D-TOPO (Invitrogen) to generate a genomic *RH12* entry clone. To generate transgenic lines expressing fluorescently tagged RH6, RH8, RH12 under their native promoters, genomic entry clones of *RH6* (as generated for the complementation of *rh6812* mutant), *RH8*, and *RH12* were transferred into the pGWB653 gateway vector^23^ by *in vitro* recombination using LR clonase (Invitrogen) to generate *gRH6-RFP*, *gRH8-RFP*, and *gRH12-RFP* constructs. The genomic entry clones of *RH8* and *RH12* were also transferred into pGWB604 and pGWB643 to generate the *gRH8-GFP* and *gRH12-CFP* constructs, respectively. All fusions were verified by sequencing before introduced into *A. tumefaciens* strain GV3101 and then transformed into different *Arabidopsis* genotypes using the floral dip method: *gRH6-RFP* was transformed into Col-0 wild-type; *gRH8-RFP* and *gRH8-GFP* were transformed into *rh8-1*; *gRH12-RFP* and *gRH12-CFP* were transformed into *rh12-2*. Transgenic T1 seeds were harvested and selected on 0.5x MS medium supplemented with 0.5% sucrose and 10 μg/mL ammonium glufosinate (Basta, Sigma).

Transgenic *Arabidopsis* lines coexpressing individual RFP-tagged RHs and DCP2-GFP were generated by transformation of *gRH6-RFP*, *gRH8-RFP*, and *gRH12-RFP* constructs into the *proDCP2:DCP2-GFP* genotype. Lines coexpressing RH6-RFP and RH8-GFP as well as RH12-RFP and RH8-GFP were generated by crossing of T2 *gRH6-RFP* and *gRH12-RFP* lines with the *gRH8-GFP* genotype. Lines coexpressing RH6-RFP and RH12-CFP was generated by crossing of T2 *gRH6-RFP* with the *gRH12-CFP* genotype. Transgenic lines coexpressing RH6-RFP and VCS-GFP as well as RH12-RFP and VCS-GFP were generated by crossing of T2 *gRH6-RFP* and *gRH12-RFP* lines with the *proVCS:VCS-GFP* genotype, whereas transgenic lines coexpressing RH8-RFP and VCS-GFP were generated by transformation of *gRH8-RFP* construct into the *proVCS:VCS-GFP* genotype. Transgenic lines coexpressing RH6-RFP and UBP1C-GFP as well as RH12-RFP and UBP1C-GFP were generated by crossing of T2 *gRH6-RFP* and *gRH12-RFP* lines with the *pro35S:UBP1C-GFP* genotype, whereas transgenic lines coexpressing RH8-RFP and UBP1C-GFP were generated by transformation of *gRH8-RFP* construct into the *gUBP1C-GFP* genotype. Generation of *rh6812 proDCP2:DCP2-GFP*, *rh6812 proVCS:VCS-GFP*, and *rh6812 pro35S:UBP1C-GFP* genotypes was performed by crossing of *proDCP2:DCP2-GFP*, *proVCS:VCS-GFP*, and *pro35S:UBP1C-GFP* genotypes with *rh6*^(+/-)^*812* plants.

#### Confocal microscopy

Confocal microscopy images were collected using a Leica SP5 laser scanning confocal microscope (Leica Microsystems) and processed with Fiji software^13^. Subcellular localization of fluorescent tagged proteins in transgenic *Arabidopsis* was performed in 4-day-old seedlings grown in a vertical position on solid 0.5x MS medium supplemented with 0.5% (w/v) sucrose. RH6-RFP, RH8-RFP and RH12-RFP fusion proteins within the root tissues were imaged using a 543 nm excitation wavelength with 25% Ar laser power, and RFP emission was collected at 608-672 nm. Visualization of the nucleus was accomplished by counter-staining of the root with 1 μg/mL 4’,6-diamidino-2-phenylindole (DAPI). For DAPI fluorescence detection, excitation light was provided by a 405 nm laser line at 5% power, and 440-540 nm fluorescent light was collected as DAPI signal. Cycloheximide, an inhibitor of translation elongation that traps mRNAs on polysomes and thereby blocks mRNP assembly^10^, was applied by transfer of seedlings into a 1.5 mL tube containing 1 mL of 200 ng/µl cycloheximide or 0.001% (v/v) DMSO (for mock control treatment) and application of vacuum infiltration for 3 min. Visualization of protein localization under a microscope was performed within minutes after infiltration. All colocalization experiments were performed using a sequential scan mode, beginning with the detection of protein with longer excitation wavelength. Colocalization analysis of RH6-RFP and RH12-CFP was performed with excitation of RFP and CFP by 543 and 458 nm laser lines at 20% power, and collection of fluorescent emission at 608-672 nm and 465-538, respectively. Colocalization analysis between RFP and GFP fusion proteins was performed using 543 nm and 488 nm laser lines at 20% power for RFP and GFP excitation, respectively. RFP and GFP emissions were collected at 577-672 nm and 494-541 nm, respectively. For comparison of DCP2-GFP, VCS-GFP, and UBP1C-GFP localization in the triple mutant *rh6812* and the wild-type, germination of *rh6812* seeds was started 12 h before those of the wild-type to compensate for their overall delayed germination. For GFP detection, a 488 nm laser at 20-35% power was used for excitation. GFP emission was captured at 494-541 nm.

#### Translating ribosome affinity purification (TRAP)

TRAP of polysomes was performed as described previously^24^. Briefly, 1 mL of pulverized tissue was added to a 15 mL tube containing 5 mL of polysome extraction buffer (PEB: 200 mM Tris, pH 9.0, 200 mM KCl, 25 mM EGTA, 35 mM MgCl_2_, 1% PTE, 1 mM DTT, 1 mM PMSF, 100 μg/mL cycloheximide, 50 μg/mL chloramphenicol) and 1% detergent mix [20% (w/v) polyoxyethylene(23)lauryl ether, 20% (v/v) Triton X-100, 20% (v/v) Octylphenyl-polyethylene glycol, 20% (v/v) Polyoxyethylene sorbitan monolaurate 20] and incubated until thawed on ice, before homogenized with a glass homogenizer. The homogenized mixture was incubated on ice for 10 min, and centrifuged at 16,000*g* at 4°C for 15 min. After centrifugation, the supernatant was transferred to a new tube and filtered through sterilized Miracloth (Millipore) to produce a clarified extract.

The clarified extract was added to Anti-FLAG M2 Protein G Dynabeads (1.5 mL) that was pre-washed twice with wash buffer [WB; 200 mM Tris (pH 9.0), 200 mM KCl, 25 mM EGTA, 35 mM MgCl_2_, 1 mM DTT, 1 mM PMSF, 100 μg/mL cycloheximide, 50 μg/mL chloramphenicol]. Binding of FLAG epitope-tagged ribosome-mRNA complexes was accomplished by incubation at 4°C for 2 h with gentle rocking. The beads were magnetically captured, the supernatant was removed, and the beads were gently resuspended in 6 mL WB for 2 min at 4°C with rocking. This step was repeated two additional times, before the beads were resuspended in 1 mL WB. The beads were washed two additional times with 1 mL WB, and the supernatant was removed. The beads were stored at −80°C until used.

#### Poly(A)^+^ RNA affinity purification

Purification of Poly(A)^+^ RNA was carried out as described previously^25^. Briefly, total and TRAP purified RNA were resuspended in 200 μL of Lysis binding buffer (LBB: 100 mM Tris-HCl, pH 8.0, 1000 mM LiCl, 10 mM EDTA, 1% (w/v) SDS, 5 mM DTT, 1.5% (v/v) Antifoam A, 5 μL/mL 2-mercaptoethanol) followed by vortexing for 5 min. Samples were incubated for 10 min at room temperature, followed by centrifugation at 13,000*g* for 10 min at room temperature. The supernatant was transferred to a new tube, and 1 μL of 12.5 μM biotin-20nt-dT oligos (Integrated DNA Technologies) was added and incubated at room temperature for 10 min. In a separate tube, 10 μL of magnetic streptavidin beads (New England Biolabs) were washed with 200 μL LBB. The sample was added to the washed beads and incubated at room temperature for 10 min with agitation. The beads were magnetically collected and washed with 200 μL of wash buffer A (WBA:10 mM Tris-HCl, pH 8.0, 150 mM LiCl, 1 mM EDTA, 0.1% (w/v) SDS), followed by wash buffer B (WBB:10 mM Tris-HCl, pH 8.0, 150 mM LiCl, 1 mM EDTA), and low-salt buffer (LSB: 20 mM Tris, pH 8.0, 150 mM NaCl, 1 mM EDTA). Subsequently, the pellet was resuspended in 10 μL of 10 mM Tris, pH 8.0 containing 1 mM 2-Mercaptoethanol, and heated at 80°C for 2 min. The beads were magnetically separated, and the supernatant was quickly transferred to a new tube. The poly(A)^+^ RNA selection process was repeated another time, and the samples were combined in a new tube before storage at −80°C.

#### Library preparation, RNA-seq data processing, and differential gene expression analysis

Non-strand specific RNA-seq libraries were prepared as described previously^25, 26^. Briefly, the purified poly(A)^+^ RNA (∼50 ng) was used for fragmentation and cDNA priming by incubating with 1.5 μL 5x RT buffer (Thermo Scientific) and 0.5 μL of 3 μg/μL random hexamer primers (Invitrogen) in a final reaction volume of 10 μL at 94 °C for 1.5 min followed by 4°C for 5 min. The fragmented RNA was used as the template for first strand cDNA synthesis by the addition of 1.5 μL 5x RT buffer, 1.5 μL of 0.1 M DTT (Invitrogen), 0.25 μL water, 0.5 μL RevertAid RT (Thermo Scientific), and 1.25 μL 10 mM dNTPs (Promega). This synthesis step was performed by incubation at 25, 42, 50 and 70°C for 10, 50, 10 and 10 min, respectively. Second strand synthesis, end preparation, and A-tailing of the first strand cDNA (15 μL) were performed in a reaction containing 1.4 μL end repair buffer (New England Biolabs), 1 μL DNA Polymerase I (Enzymatics), 0.1 μL RNaseH (Enzymatics), 0.4 μL T4 Pol + PNK mix (end repair module, New England Biolabs), 0.2 μL Taq DNA polymerase (GBioscience), 1 μL 10 mM dNTPs (Promega), and 0.9 μL water with the following program: 16°C for 20 min, 20°C for 20 min, and 72°C for 20 min. The cDNA was purified using 30 μL AMPure XP beads (Beckman), washed twice with 120 μL 80% ethanol, and used in the adaptor ligation step. Adaptor ligation was accomplished using 5 μL of 2x Rapid T4 ligase buffer (Enzymatics), 3 μL 1 μM annealed universal Y-shaped adapter, 0.25 μL T4 DNA Ligase HC (Enzymatics), 1.75 μL water with incubation at room temperature for 15 min. Next, 25 μL of Ampure XP Bead Resuspension buffer (ABR: 15% PEG 8000, 2.5M NaCL) was added to the reaction and incubated for 5 min at room temperature. The beads were magnetically separated, washed twice with 200 μL 80% ethanol, and resuspended in 10 μL water. Following incubation for 2 min at room temperature, the beads were magnetically separated, and the supernatant was transferred to a new tube. Thia adapter-coupled cDNA (10 μL) was used for PCR enrichment and adapter extension in a reaction containing 4 μL 5X Kapa HiFi Fidelity buffer (Kapa Biosystems), 1 μL of 2 μM indexed primer, 1 μL of 2 μM PE1 primer, 1 μL of 8 μM EnrichS1 and EnrichS2 primers, 0.2 μL Kapa HiFi DNA polymerase (Kapa Biosystems), 0.5 μL of 10 mM dNTPs (Promega), and 2.3 μL water. The PCR reactions included denaturation by incubation at 98°C for 30 sec, followed by 15 cycles of denaturation at 98°C for 10 sec, annealing at 65°C for 30 sec, extension at 72°C for 30 sec, and a final extension at 72°C for 5 min. The amplified cDNA libraries of 200-300 bp were purified using acrylamide gel electrophoresis and elution. Library yield and concentration were checked by Qubit 2.0 Fluorometer (Life Technologies) and Bioanalyzer. The libraries were multiplexed and sequenced on short-read Illumina sequencers at the UC Riverside Institute for Integrative Genome Biology Genomics Core facility.

Analysis of RNA-seq data was performed on a Linux cluster using a combination of command-line software and R packages including the RNA-Seq data analysis pipeline package systemPipeR^27^. Multiplexed libraries were demultiplexed using their unique barcodes. Reads were trimmed to correct for adapter contamination with standard settings. Bowtie 2 (v2.2.5) was used to build the TAIR10 reference genome from individual chromosome Fasta files downloaded from www.araport.org. Reads were aligned to this assembled *Arabidopsis* genome using Bowtie2 (v2.2.5) /Tophat (v2.0.14). Aligned reads were counted against the Araport11 genome release 201606 using the summarizeOverlaps function^28^ in “union” mode. Differential gene expression analysis was performed using the generalized linear model within the edgeR package^29^. Genes that contained ≤ 2 counts per million (CPM) in at least two samples were removed. Genes which passed the first filter were passed through a second filter that retained only those with RPKM > 5 in at least two samples. The calculated *p* value for each gene was corrected by applying the Benjamini-Hochberg method, to provide a false discovery rate (FDR). Genes with fold change > |2| and FDR < 0.01 were considered differentially expressed. GO analysis was conducted in R using the systemPipeR package environment^27^.

#### Global mRNA decay analysis

Col-0 wild-type and *rh6*^(+/-)^*812* plated on sterile solid medium, and stratified for 2 days were grown vertically at 22°C in a Conviron growth chamber under ∼75 μmol photons·s^-1^·m^-2^ constant illumination for 5 days. Seedlings with clear triple *rh6812* mutant phenotypes (∼12-15%) were placed on fresh growth media before cordycepin treatment. Col-0 seedlings were similarly handled to standardize the treatment. The experiment was performed in 3 independent biological replicates with ∼100 *rh6812* seedlings and ∼50 Col-0 seedlings per sample. Cordycepin treatment was carried out as described^30^. Briefly, seedlings were preincubated in 3 mL of incubation buffer (15 mM sucrose, 1 mM KCl, 1 mM PIPES pH 6.25, 1 mM sodium citrate) with rotation at 100 rpm in 35 mm x 10 mm Petri dishes for 15 min prior to replacement of the bathing solution with 3 mL of fresh buffer containing 1 mM cordycepin (Chengdu Biopurify Phytochemicals), followed by vacuum infiltration for 1 minute. T_0_ samples were harvested and frozen immediately, with the remaining samples subjected immediately to two additional vacuum infiltration of 1 min-vacuum with 1 min-decompression intervals. Tissues were collected 15, 30, 60, and 120 min following the first vacuum infiltration, frozen in liquid nitrogen, and stored at −80°C.

Total RNA was isolated from pulverized tissues by addition of 500 μL PureLink™ Plant RNA Reagent (Invitrogen). The samples were vortexed and incubated at room temperature for 5 min, followed by centrifugation at 12,000*g* for 2 min at room temperature. The supernatant was transferred to a tube containing 5 M NaCl (equal volume). Chloroform (300 μL) was added and the sample was mixed thoroughly before centrifugation at 12,000*g* for 10 min at 4°C, and transfer of the upper aqueous phase to a new tube. RNA was precipitated by addition of isopropanol (equal volume), thorough mix, incubation at room temperature for 10 min, and pelleted by centrifugation at 12,000*g* for 10 min at 4°C. The supernatant was removed, and the pellet was washed with 1 mL 70% (v/v) ethanol and resuspended in 30 μL RNAsecure™ Resuspension Solution (Invitrogen), and DNase I-digested with 1 μL of TURBO DNase (Invitrogen) for 15 min at 37°C. The RNA was precipitated in 0.1 volume of 5 M ammonium acetate and 2.5 volumes of ethanol, and pelleted by centrifugation at 12,000*g* for 20 min at 4°C. DNase-treated RNA was resuspended in RNase-free water, and submitted to the University of Utah Genomics core for RNA-seq library preparation and sequencing. RNA-seq libraries were constructed using the Illumina TruSeq Total RNA Sample Prep Kit with Ribo-Zero (Plant) (Illumina). A total of 30 libraries was multiplexed so that all time course and genotype samples in each biological replicate (10 samples) were loaded onto a single lane.

Processing of RNA-seq data was performed using the package systemPipeR^27^, with the same settings as in the transcriptome-translatome analysis. Data normalization as well as modeling of mRNA decay and genotype effect were performed using the Bioconductor RNAdecay package^31^. Genotypic effect on mRNA decay for each transcript was fitted to 6 possible exponential decay models with α as an estimated parameter of the initial primary decay rate and β as an estimated parameter of the secondary decay of the α decay rate, using likelihood functions. The model with the lowest Akaike information criterion (AICc) was selected for each transcript. Only the model with the estimated parameter of variance (σ^2^) less than 0.0625 was accepted for a downstream analysis. mRNA half-life was calculated as *t*_1/2_ = ln(2)/α. Comparison of RH and VCS substrates was limited to mRNAs with a *t*_1/2_ in Col-0 of less than 12 h (*n* = 12,697) as the maximum *t*_1/2_ of RH substrates was 11.5 h.

#### SA quantification

Quantification of SA in was carried out by liquid chromatography–mass spectrometry (LC-MS), using deuterated SA as internal standard. Extraction and quantification were performed as previously described^32^.

#### Pathogen infection and tissue staining

*Hyaloperonospora arabidopsidis* (*Hpa*) Noco2 was grown and propagated as described previously^33^. Arabidopsis seeds were grown on MS medium and 4-day-old seedlings were transferred to soil. Seven-day-old seedlings were spray-infected with *Hpa* spores at a concentration of 5 x 10^4^ spores per ml of water with Preval sprayers (http://www.prevalspraygun.com). Infected plants were kept in a humid growth chamber (12 h light/12 h dark, 18°C) and scored for *Hpa* growth 7 days post infection by counting spores using a hemocytometer to determine the spore density of a suspension of 30 infected seedlings per 1 ml of water. Trypan blue staining was performed as described previously^33^.

#### Statistical analysis

Statistical analysis was performed in R software using agricolae package^34^.

## Supplementary Figures

**Supplementary Fig.1.**
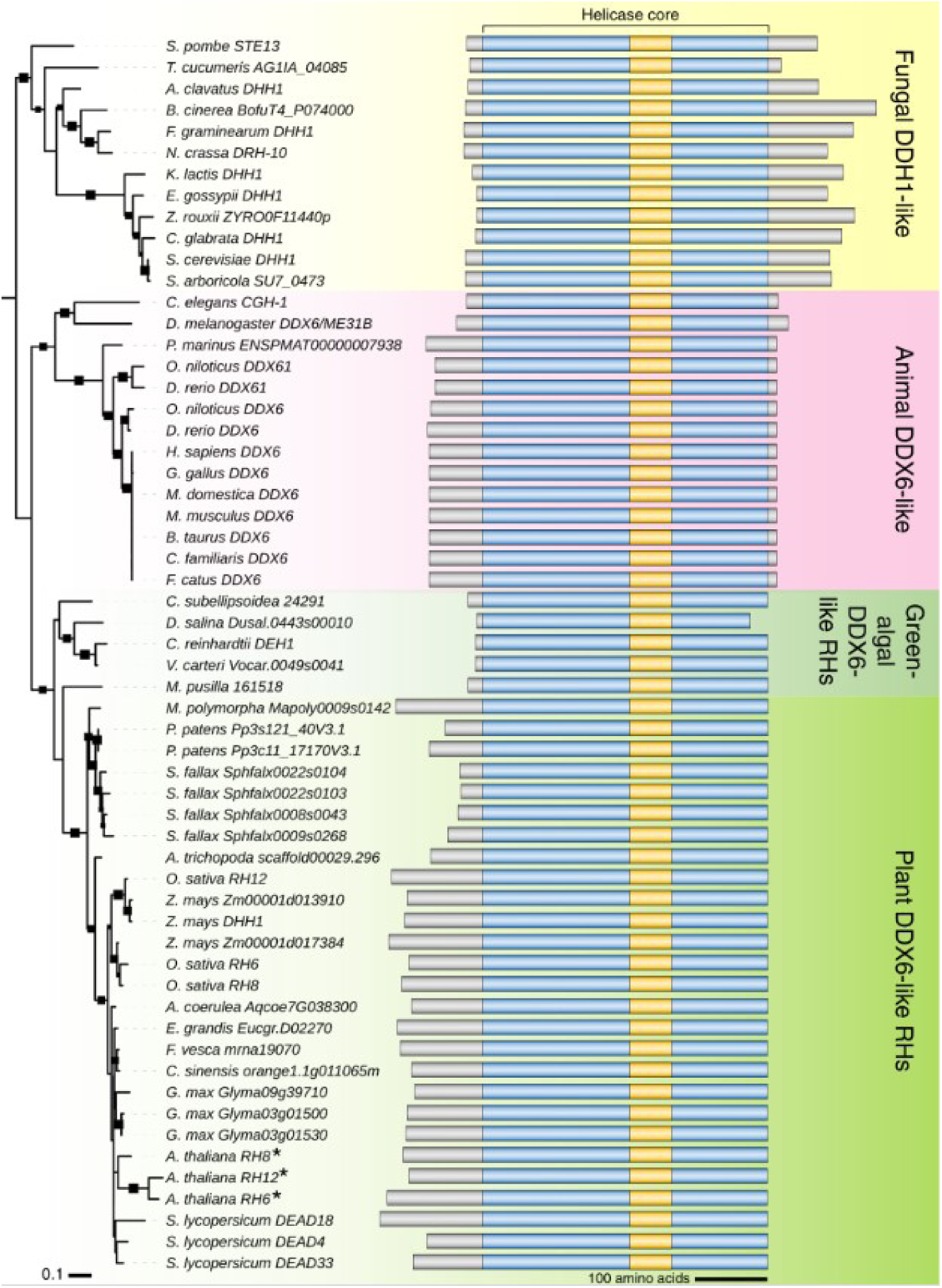
Phylogenetic relationship and schematic representation of primary sequence structure of eukaryotic DHH1/DDX6-like DEAD-box RNA helicases. Evolutionary history was inferred from 58 representative DHH1/DDX6-like helicases across different eukaryotic lineages. The sequences were aligned using the Muscle algorithm and the phylogenetic tree was generated with the Maximum Likelihood method with 1000 bootstrap replicates. Evolutionary rate differences among sites were modelled with a discrete Gamma distribution. The tree is shown to scale, with branch lengths measured as the number of substitutions per site. Black rectangular boxes on the tree branches depict bootstrap values greater than 50%, with the size of the boxes being proportional to the bootstrap values. Grey boxes represent the N- and C-terminal extensions; blue boxes, RecA-like domains; yellow boxes, linker regions.

**Supplementary Fig.2.**
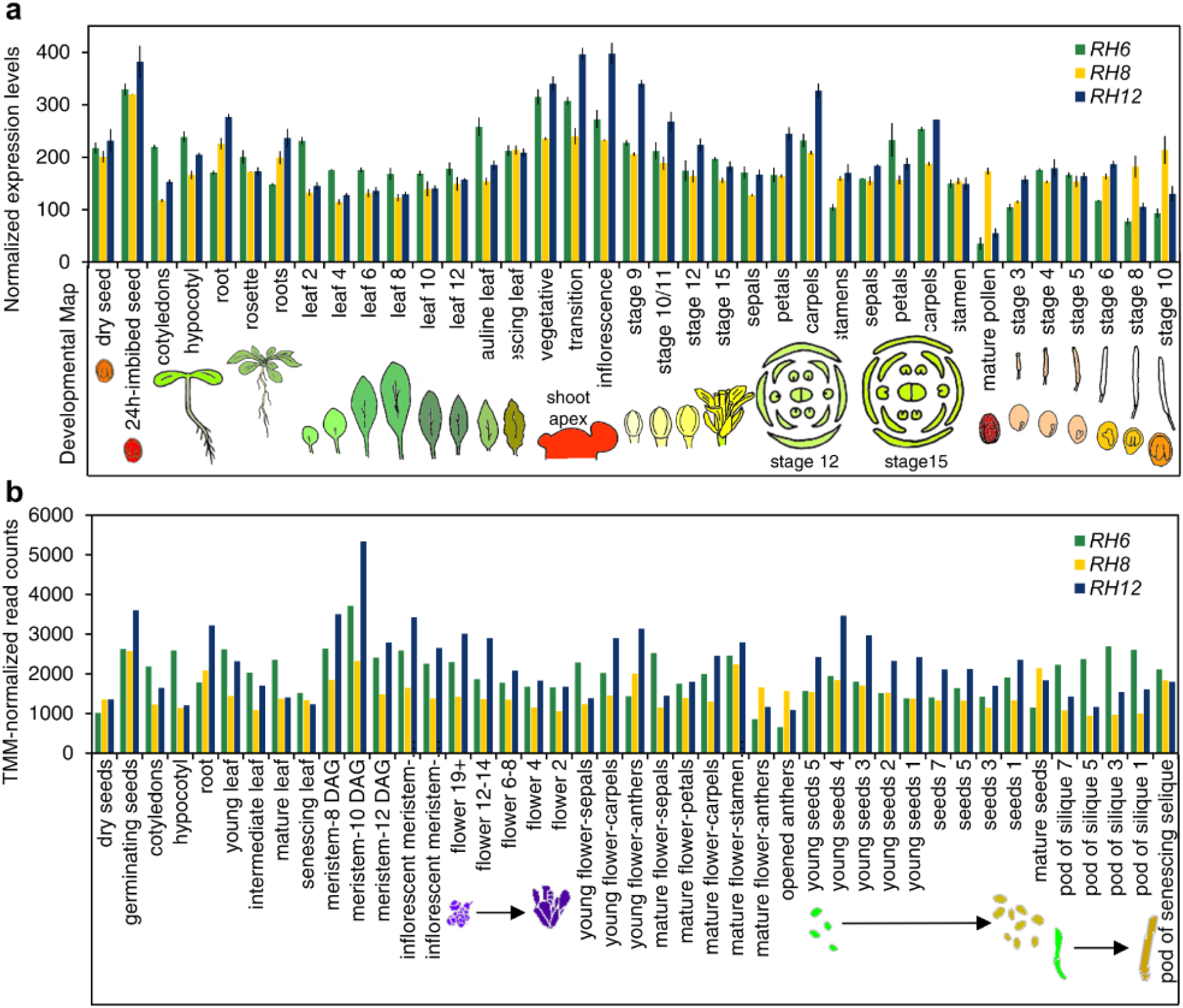
Tissue-specific expression profiles of *RH6*, *RH8* and *RH12* transcripts. **a,b**, Expression pattern of *RH6*, *RH8*, and *RH12* across different organs and developmental stages based on the publicly available collection of microarray (**a**) and RNA-seq (**b**) data. Microarray and RNA-seq data were extracted from the *Arabidopsis* eFP browser (http://bar.utoronto.ca/efp/cgi-bin/efpWeb.cgi) and TraVa database (http://travadb.org/), respectively.

**Supplementary Fig.3.**
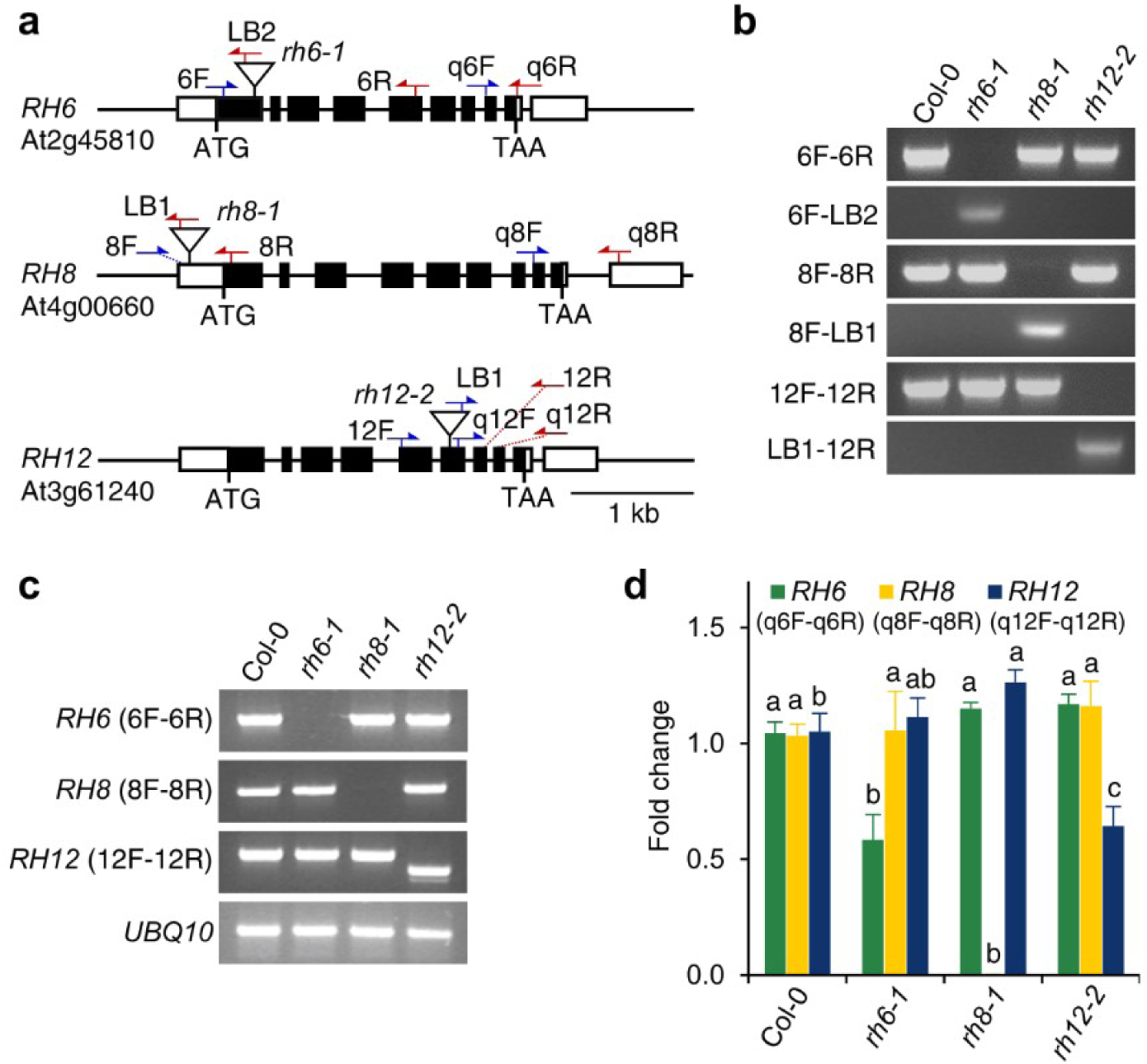
Characterization of Arabidopsis *rh6*, *rh8* and *rh12* mutants. **a**, Schematic representation of the *RH6*, *RH8* and *RH12* loci and their associated T-DNA insertion alleles. Untranslated and coding regions are depicted as white and black boxes, respectively. Introns are depicted as lines. “ATG” and “TAA” indicate the translational initiation and stop codon, respectively. Inverted white triangles refer to the positions of T-DNA insertions in the *rh6-1*, *rh8-1* and *rh12-2* alleles. Arrows indicate the orientations and approximate positions of primers used for molecular genotyping of the genes or the T-DNAs: blue, forward primers; red, reverse primers. **b**, PCR-based genotyping of the homozygous *rh6-1*, *rh8-1* and rh*12-2* alleles. Gene and T-DNA-specific primers represented in **a** were used for PCR amplification of genomic DNA from Col-0 and homozygous *rh6-1*, *rh8-1* and *rh12-2* insertion alleles. **c**, Reverse transcriptase (RT)-PCR analysis of *RH6*, *RH8*, and *RH12* transcripts. Primers flanking the T-DNAs as indicated in **a** were used for PCR-based specific detection of *RH6*, *RH8* and *RH12* transcripts from the homozygous *rh6-1*, *rh8-1* and rh*12-2* alleles in comparison to Col-0. *UBQ10* was used as an internal control. **d**, RT-qPCR analysis of *RH6*, *RH8* and *RH12* transcript levels in 7-day-old seedlings of the homozygous *rh6-1*, *rh8-1* and *rh12-2* mutants. Gene-specific primers located downstream of the T-DNA as indicated in **a** were used. Relative transcript fold-change was calculated using *PP2AA2* as a reference. Error bars, SD (*n*=3). Statistical significance was determined by ANOVA, followed by Tukey’s Honest Significant Difference (HSD) test. Means that are significantly different from each other (*p* < 0.05) are denoted by different letters.

**Supplementary Fig.4.**
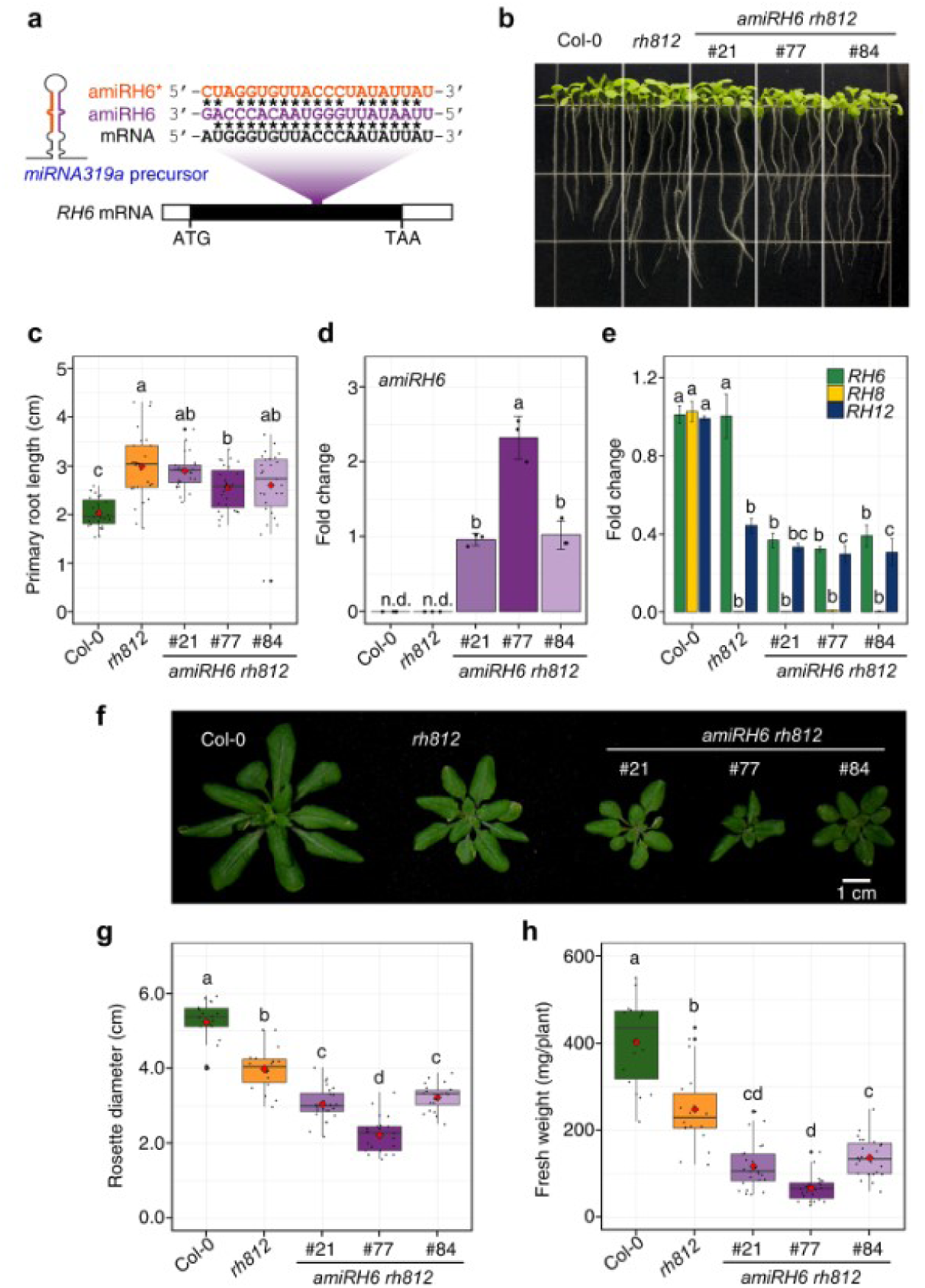
Reduced expression of *RH6*, *RH8* and *RH12* affects plant growth. **a**, Schematic diagram and sequences of a 21-nucleotide artificial miRNA and its target site on *RH6* expressed in the Arabidopsis *MIR319a* backbone under the 35S promoter. **b-c**, Phenotype and primary root lengths of 7-day-old seedlings of the wild-type Col-0, the double *rh812* mutant, and three independent *amiRH6* lines generated in *rh812* background (*amiRH6 rh812*). **d**, Levels of artificial miRNA *amiRH6* in 7-day-old seedlings of the *amiRH6 rh812* lines in **b** determined by pulsed stem-loop RT-qPCR. miRNA fold-change was calculated relative to line #21 using *U6* RNA as a reference. Error bars, SD (*n*=3); n.d., not detectable. **e**, RT-qPCR analysis of *RH6*, *RH8* and *RH12* transcript levels in 7-day-old seedlings of Col-0, *rh812* and three *amiRH6 rh812* lines. Relative transcript fold-change was calculated using *PP2AA2* as a reference. Error bars, SD (*n*=3). **f**, Rosette growth phenotype of 28-day-old plants of Col-0, *rh812* and three *amiRH6 rh812* lines. **g-h**, Rosette diameters and fresh weights (*n* = 18-24) of 28-day-old plants of the genotypes presented in **f**. Statistical significance was determined by ANOVA, followed by Tukey’s HSD test. Means that are significantly different from each other (*p* < 0.05) are denoted by different letters.

**Supplementary Fig.5.**
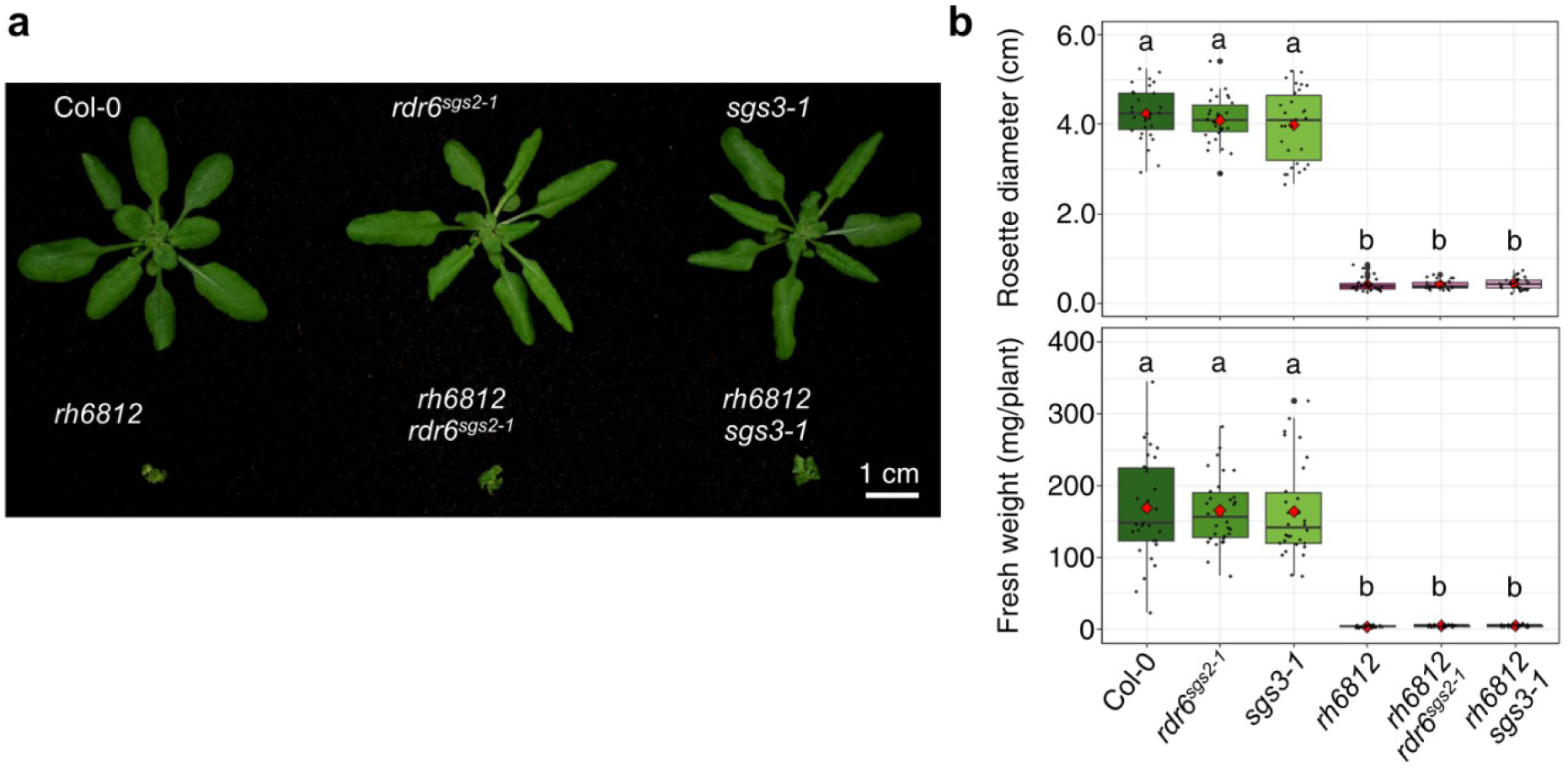
The triple *rh6812* mutant phenotype is *RNA-DEPENDENT RNA POLYMERASE 6* and *SUPPRESSOR OF GENE SILENCING 3* independent. **a**, Representative rosette growth phenotype of 28-day-old plants of Col-0, single *rdr6^sgs2-1^* and *sgs3-1* mutants, triple *rh6812* mutant, and quadruple *rh6812 rdr6^sgs2-1^* and *rh6812 sgs3-1* mutants. **b**, Rosette diameter and fresh weight of 28-day-old plants of the genotypes presented in **a**. Statistical significance was determined by ANOVA (*n* = 30), followed by Tukey’s HSD test. Data were log transformed. Means that are significantly different from one another (*p* < 0.05) are denoted by different letters.

**Supplementary Fig.6.**
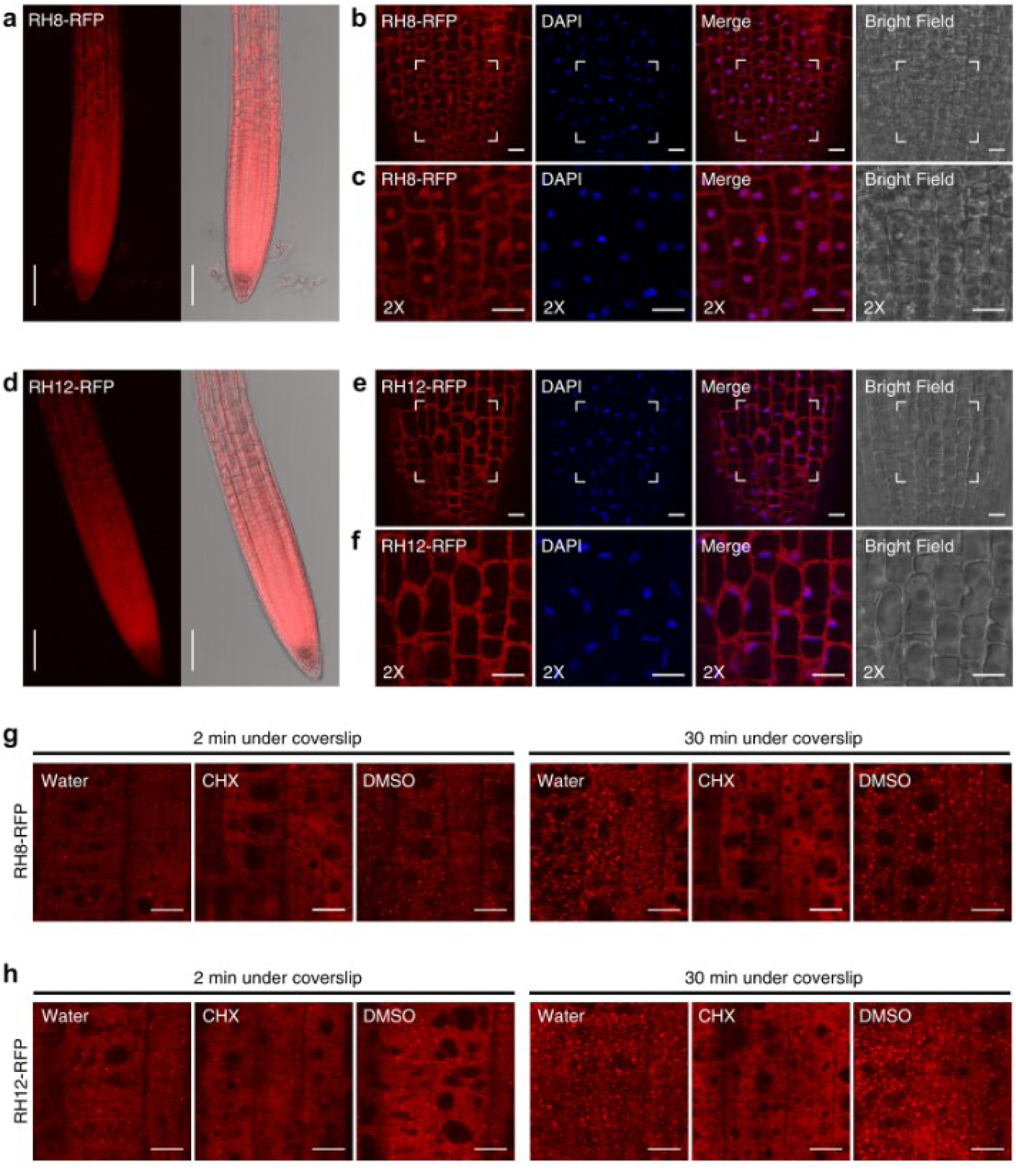
Subcellular localization of Arabidopsis RH8 and RH12. **a**, Confocal images of RFP fluorescence in root tissues of transgenic 4-day-old seedlings expressing *genomic(g)RH8-RFP* under control of its native promoter. Left panel: maximum projection over 12 *z*-planes for a total volume of 55 µm; right panel: overlay with bright field image. Bars = 100 µm. **b**, RFP fluorescence of the root meristem region of the plants described in **a** counter-stained with 4’,6-diamidino-2-phenylindole (DAPI) for nuclear visualization. **c**, Magnified images of framed areas in **b**. Bars in **b** and **c** = 10 µm. **d-f**, Confocal images of *gRH12-RFP* under control of its native promoter presented in the same manner as for *gRH8-RFP* in **a-c**. **g-h**, Confocal images of RFP fluorescence in the root meristem regions of transgenic seedlings expressing *gRH8-RFP* and *gRH12-RFP*. Four-day-old seedlings were submerged in water under a coverslip and imaged after 2 and 30 min. Seedlings were pre-treated with 0.001% (v/v) dimethyl sulfoxide (DMSO) with or without cycloheximide (CHX; 200 ng µl^-1^) for 3 min by vacuum infiltration before imaging as described. More foci are evident after vacuum infiltration of DMSO than with water-only immersion, but these inhibited by CHX. Images representative of *n* ≥ 3 seedlings per bioreplicates. Bars = 10 µm.

**Supplementary Fig.7.**
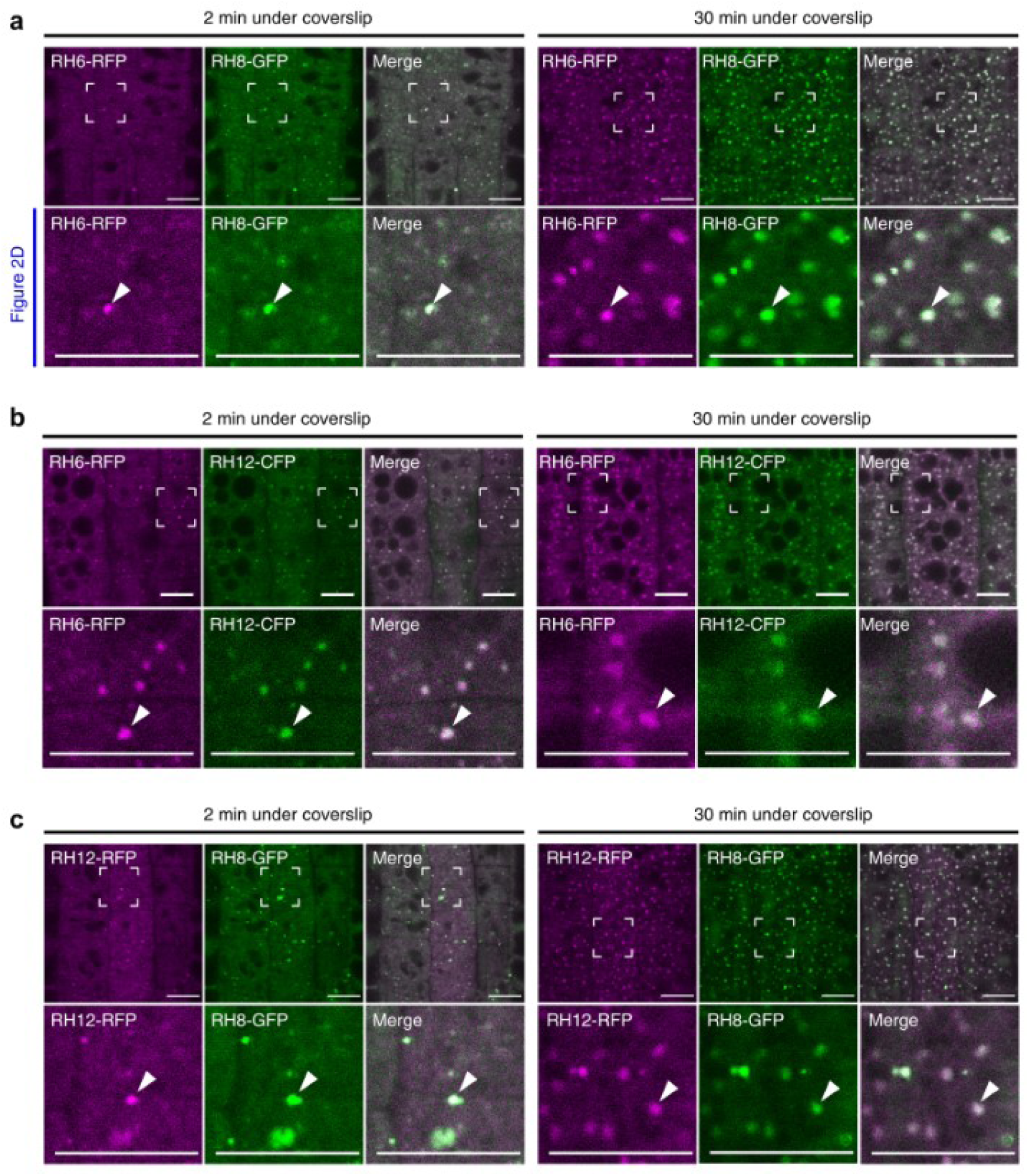
RH6, RH8 and RH12 overlap in cytoplasmic foci. **a**, Confocal images of RFP and GFP fluorescence in the root meristem of 4-day-old seedlings co-expressing *gRH6-RFP* and *gRH8-GFP*. The seedlings were submerged in water under a coverslip, and subcellular localization of RH6-RFP and RH8-GFP was determined after 2 and 30 min. RFP and GFP signals were false-colored into magenta and green, respectively. Lower panels are magnified images of framed areas in the upper panels, and are as presented in Fig. 2d. Arrowheads denote an example of foci where RH6-RFP and RH8-GFP colocalize. Bars = 10 µm. **b**, Confocal images of 4-day-old seedlings co-expressing *gRH6-RFP* and *gRH12-CFP*. **c**, Confocal images of 4-day-old seedlings co-expressing *gRH12-RFP* and *gRH8-GFP*. Data in **b** and **c** are presented the same way as in **a** except that RFP and CFP signals in **b** were false-colored into magenta and green, respectively.

**Supplementary Fig.8.**
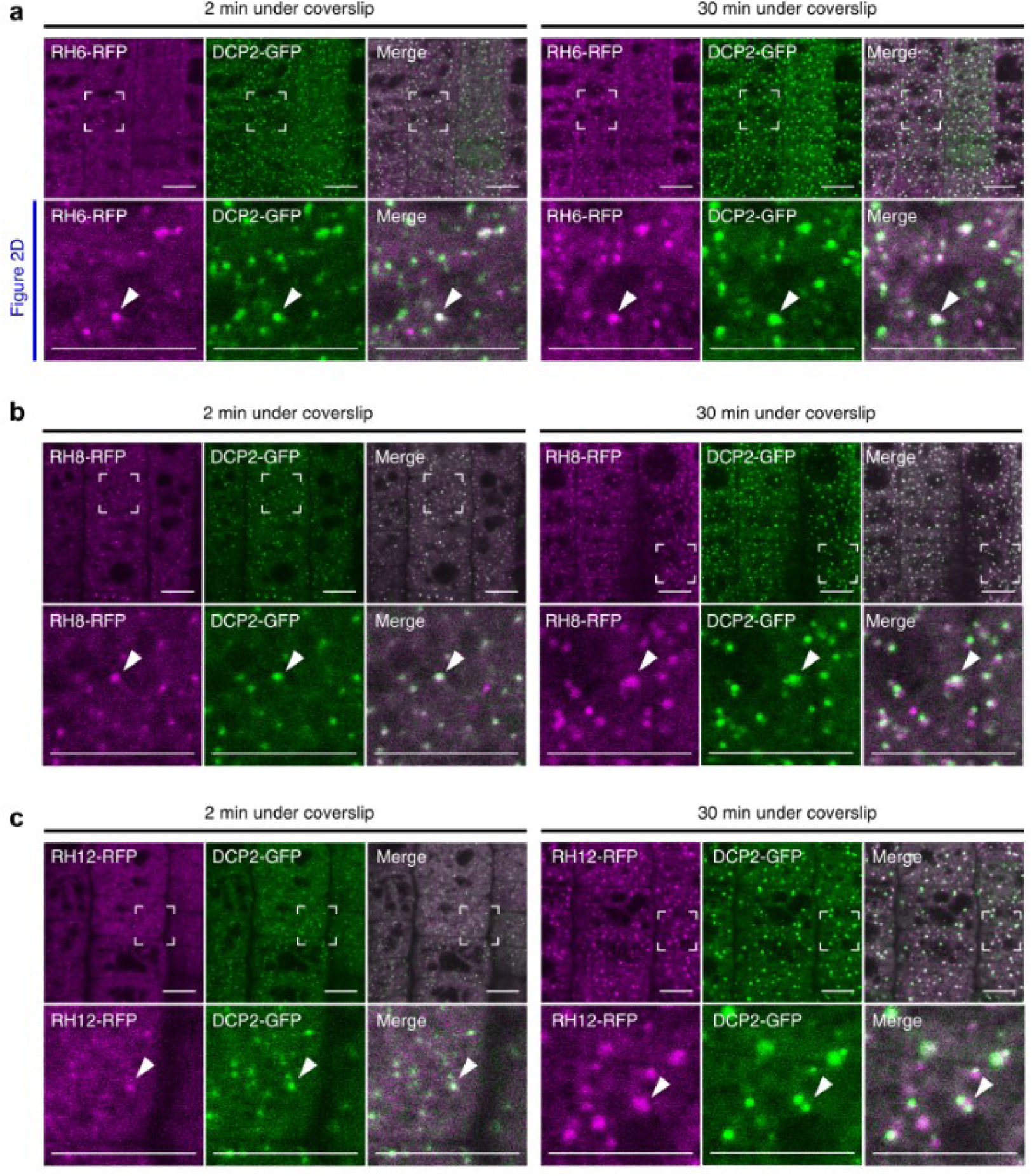
RH6, RH8 and RH12 foci overlap with those of the decapping enzyme DCP2. **a**, Confocal images of RFP and GFP fluorescence in the root meristem of 4-day-old seedlings co-expressing *gRH6-RFP* and *proDCP2:DCP2-GFP*. The seedlings were submerged in water under a coverslip, and subcellular localization of RH6-RFP and DCP2-GFP was determined after 2 and 30 min. RFP and GFP signals were false-colored into magenta and green, respectively. Lower panels are magnified images of framed areas in the upper panels and are as presented in Fig. 2d. Arrowheads denote an example of foci where RH6-RFP and DCP2-GFP colocalize. Bars = 10 µm. **b**, Confocal images of 4-day-old seedlings co-expressing *gRH8-RFP* and *proDCP2:DCP2-GFP*. **c**, Confocal images of 4-day-old seedlings co-expressing *gRH12-RFP* and *proDCP2:DCP2-GFP*. Data in **b** and **c** are presented the same way as in **a**.

**Supplementary Fig.9.**
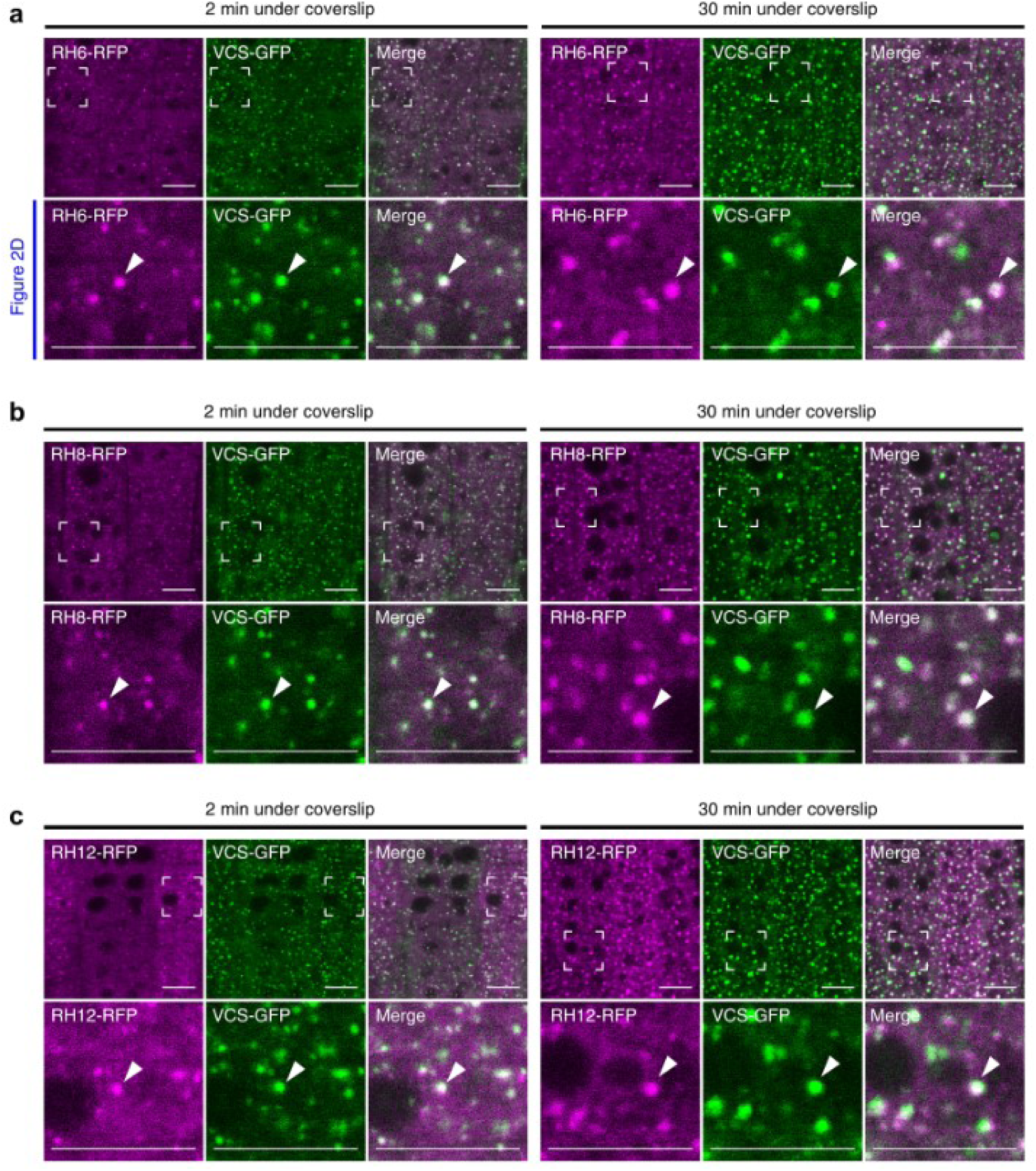
RH6, RH8 and RH12 complexes overlap with those of the core decapping protein VCS. **a**, Fluorescence microscopy images of 4-day-old seedlings co-expressing *gRH6-RFP* and *proVCS:VCS-GFP*. The seedlings were submerged in water under a coverslip, and subcellular localization of RH6-RFP and VCS-GFP was determined in the root meristem regions at 2 and 30 min of submergence. RFP and GFP signals were false-colored in magenta and green, respectively. Lower panels are magnified images of framed areas in the upper panels and are as presented in Fig. 2d. Arrowheads indicate representative foci where RH6-RFP and VCS-GFP colocalize. Bars = 10 µm. **b**, Confocal images of 4-day-old seedlings co-expressing *gRH8-RFP* and *proVCS:VCS-GFP*. **C**, Confocal images of 4-day-old seedlings co-expressing *gRH12-RFP* and *proVCS:VCS-GFP*. Data in **b** and **c** are presented the same way as in **a**.

**Supplementary Fig.10.**
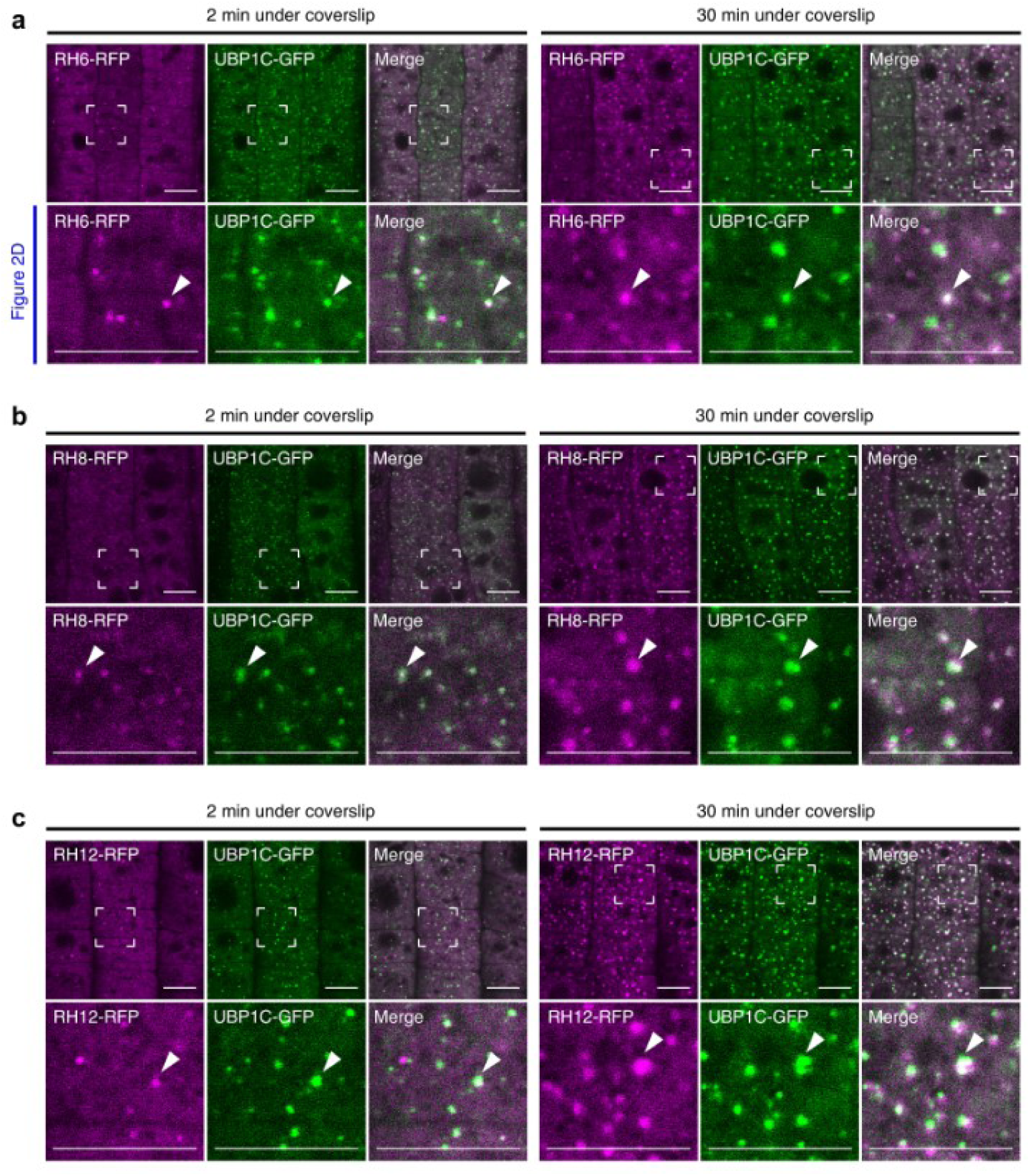
RH6, RH8 and RH12 complexes overlap with UBP1C stress granules. **a**, Confocal images of 4-day-old seedlings co-expressing *gRH6-RFP* and *pro35S:UBP1C-GFP*. Seedlings were submerged in water under a coverslip, and subcellular localization of RH6-RFP and UBP1C-GFP was determined after 2 and 30 min of submergence. RFP and GFP signals were false-colored into magenta and green, respectively. Lower panels are magnified images of framed areas in the upper panels and are as presented in Fig. 2d. Arrowheads point to a representative complex where RH6-RFP and UBP1C-GFP colocalize. Bars = 10 µm. **b**, Confocal images of 4-day-old seedlings co-expressing *gRH8-RFP* and *pro35S:UBP1C-GFP*. **c**, Confocal images of 4-day-old seedlings co-expressing *gRH12-RFP* and *pro35S:UBP1C-GFP*. Data in **b** and **c** are presented the same way as in **a**.

**Supplementary Fig.11.**
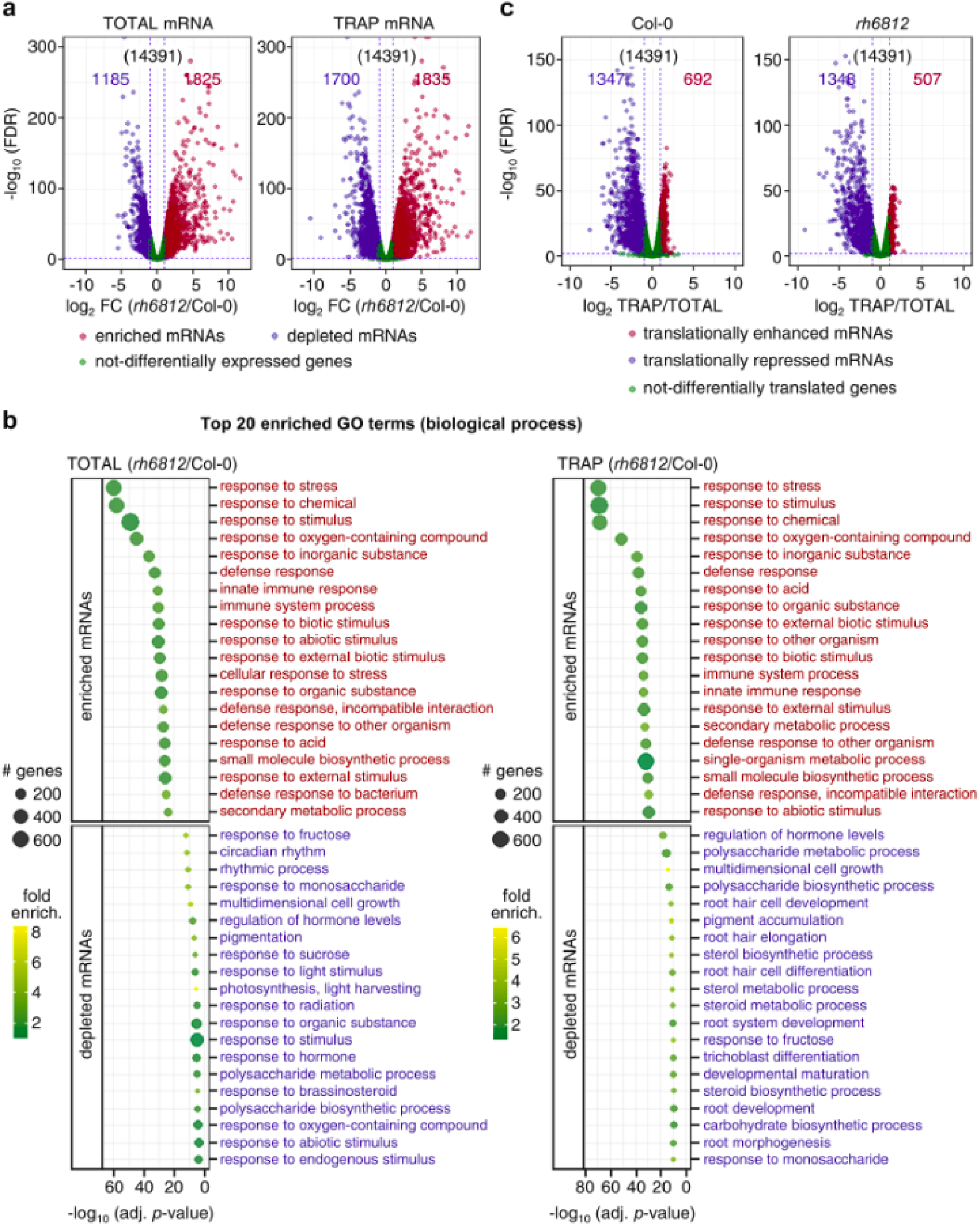
The triple *rh6812* mutant transcriptome and translatome are enriched with stress/defense-responsive mRNAs but depleted of those required for growth and development. **a**, Volcano plots of change in Total and TRAP mRNA abundance of *rh6812* relative to Col-0 based on differential abundance analysis by edgeR. The log_2_ fold-change (FC) is shown on the x-axis, and the negative log_10_ of the false-discovery rate (FDR) is shown on the y-axis. Genes with |log_2_ FC| ≥ 1 and FDR < 0.01 were considered differentially expressed. Total number of genes in the analysis is given in parentheses. **b**, GO functional categories (biological process) of gene transcripts enriched or depleted in *rh6812* relative to Col-0 as identified in **a**. Twenty non-redundant terms with the lowest adjusted *p*-values are presented. Fold enrichment (fold enrich.) represents the number of genes observed relative to the expected number in each category. **c**, Volcano plots of change in translational status calculated by comparison of steady-state Total and TRAP mRNA abundance in Col-0 and *rh6812* based on differential abundance analysis by edgeR. The log_2_ FC is shown on the x-axis, and the negative log_10_ of the FDR is shown on the *y*-axis. Genes with |log_2_ FC| ≥ 1, FDR < 0.01 were considered translationally enhanced or repressed. Total number of genes used in the analysis is presented in parentheses.

**Supplementary Fig.12.**
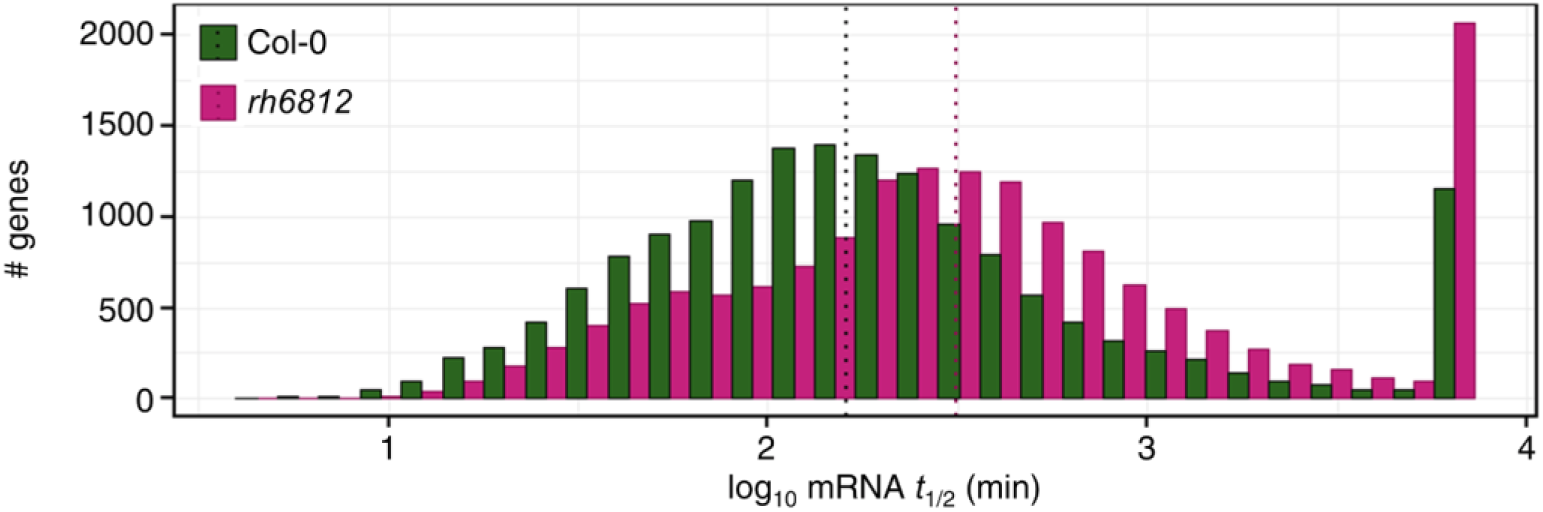
Histograms of mRNA half-life (*t*_1/2_) distributions in the Col-0 wild-type and the triple *rh6812* mutant. The analysis was performed on 16,025 genes for both genotypes representing a high-quality decay dataset (σ^2^ < 0.0625) from a total of 18092 genes. Vertical black and pink lines represent the median mRNA *t*_1/2_ in Col-0 and *rh6812*, respectively.

**Supplementary Fig.13.**
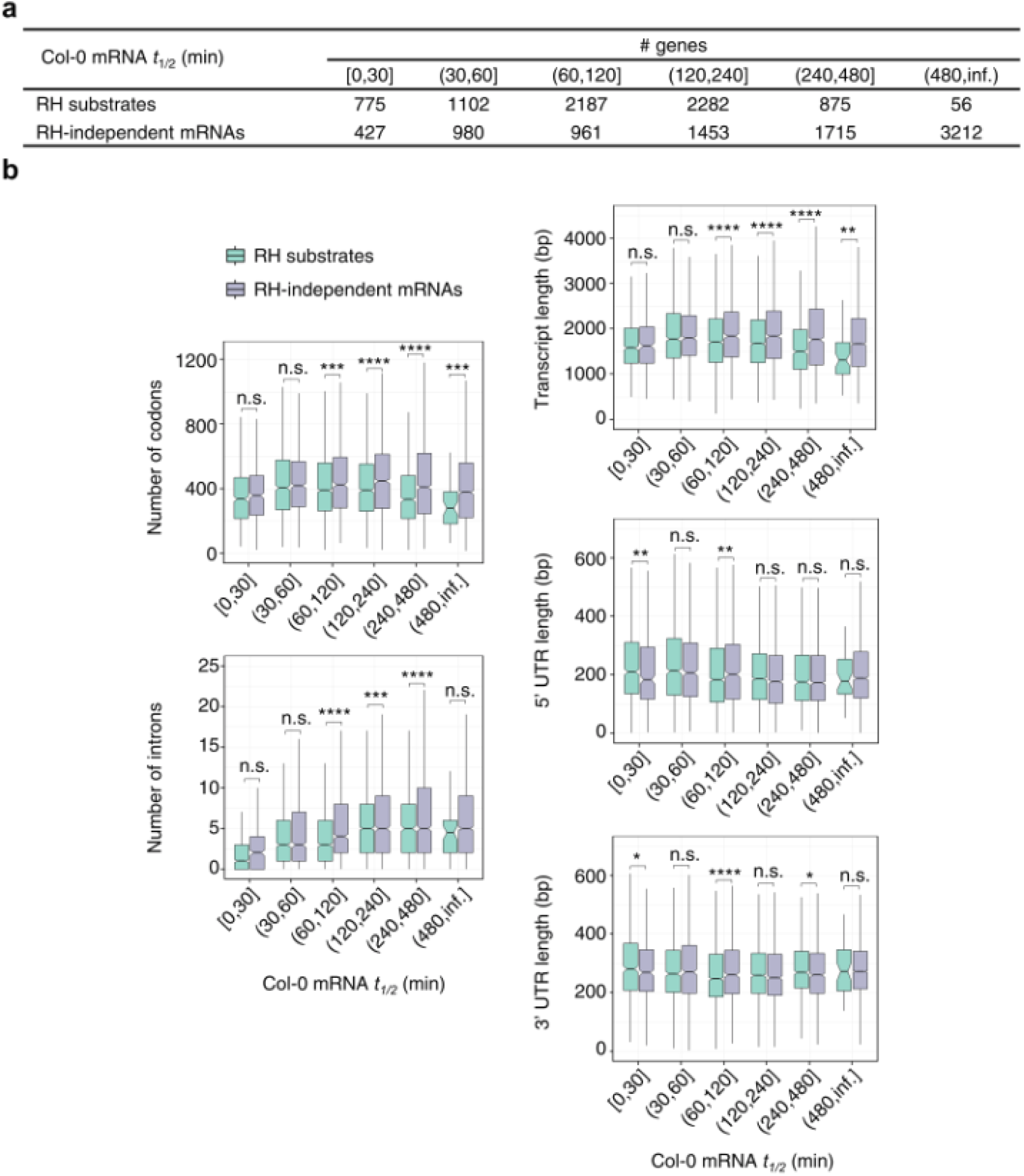
Characteristics of RH substrates. **a**, Number of genes encoding RH substrates and RH-independent mRNAs binned into six classes based on mRNA *t*_1/2_ in Col-0. **b**, Box plots comparing mRNA features between genes that encode RH substrates and RH-independent mRNAs. * denotes *p*-value < 0.05; ** denotes *p*-value < 0.01; *** denotes *p*-value < 0.001; **** denotes *p*-value < 0.0001; n.s., not significantly different (Wilcoxon test).

**Supplementary Fig.14.**
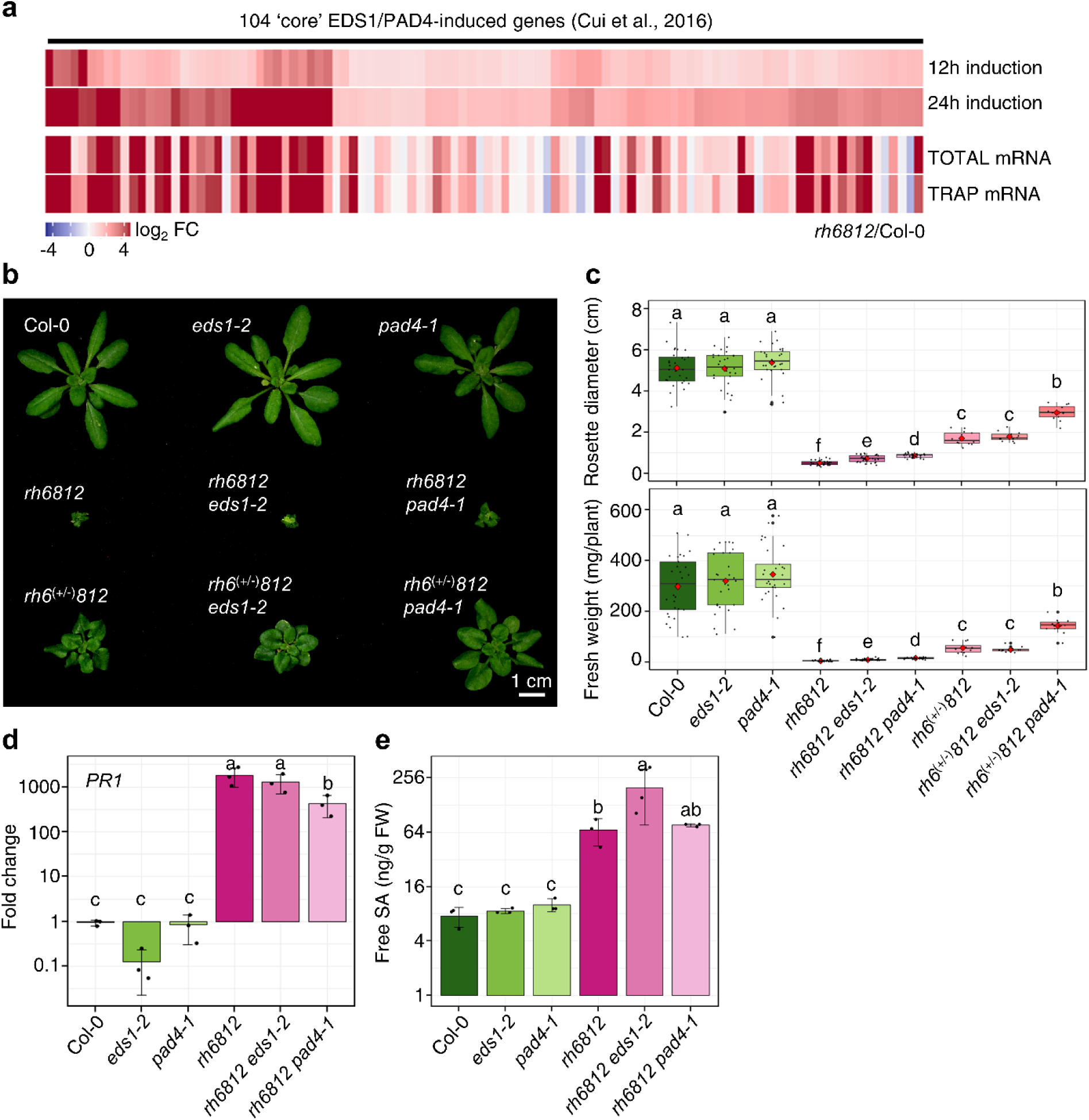
The *rh6812* mutant exhibits autoimmunity in a PAD4-partially dependent but EDS1-independent manner. **a**, Heatmap representing relative fold change in Total and TRAP mRNA abundance for 104 ‘core’ EDS1/PAD-induced genes^35^ in *rh6812* relative to Col-0. Individual genes are presented as column with the upper two rows showing their log_2_ FC following 12 and 24 hours of EDS1 and PAD4 overexpression (Cui et al., 2016), whereas the lower two rows representing their log_2_ FC in Total and TRAP populations of *rh6812* relative to Col-0. **b**, Representative rosette growth phenotype of 28-day-old plants of Col-0, *eds1-2*, *pad4-1*, *rh6812*, *rh6812 eds1-2*, *rh6812 pad4-1*, *rh6*^(+/-)^*812*, *rh6*^(+/-)^*812 eds1-2* and *rh6*^(+/-)^*812 pad4-1* genotypes. **c**, Rosette diameter and fresh weight of 28-day-old plants (*n*=12-30) of the genotypes presented in **b**. **d**, RT-qPCR analysis of *PR1* transcripts in 7-day-old seedlings of the *rh6812 eds1-2* and *rh6812 pad4-1* genotypes relative to *rh6812* and the control genotypes Col-0, *eds1-2* and *pad4-1*. Relative transcript fold-change was calculated using *ACT1* as a reference. Error bars, SD (*n*=3). **e**, Comparison of SA levels in 7-day-old seedlings of the genotypes as in **d**. Error bars, SD (*n*=3). Statistical significance in **c-e** was determined by ANOVA followed by Tukey’s HSD test. Data in **c** and **d** were log transformed. Significantly difference of means (*p* < 0.05) are represented by different letters.

